# Post-senescence reproductive rebound in *Daphnia* associated with reversal of age-related transcriptional changes

**DOI:** 10.1101/2024.04.13.589373

**Authors:** Ishaan Dua, A. Catherine Pearson, Rachael L. Lowman, Leonid Peshkin, Lev Y. Yampolsky

## Abstract

A long-lived species of zooplankton microcrustaceans, *Daphnia magna,* sometimes exhibits late-life rebound of reproduction, briefly reversing reproductive senescence. Such events are often interpreted as terminal investment in anticipation of imminent mortality. We demonstrate that such post-senescence reproductive events (PSREs) neither cause not anticipate increased mortality. We analyze an RNAseq experiment comparing young, old reproductively senescent, and old PSRE *Daphnia* females. We first show that overall age-related transcriptional changes are dominated by the increase transcription of guanidine monophosphate synthases and guanylate cyclases, as well as two groups of presumed transposon-encoded proteins and by a drop in transcription of protein synthesis-related genes. We then focus on gene families and functional groups in which full or partial reversal of age-related transcriptional changes occur. This analysis reveals reversal, in the PSRE individuals, of age-related up-regulation of apolipoproteins D, lysosomal lipases, and peptidases and of age-related down-regulation of E3 ubiquitin kinases, V-type proton ATPases, and numerous proteins related to mitochondrial and muscle functions. While it is not certain which of these changes enable reproductive rejuvenation, and which are by-products of processes that lead to it, we present some evidence that post-senescence reproductive events are associated with the reversal of age-related protein and lipid aggregates removal, apoptosis, and with restoration of mitochondrial integrity.

**Significance statement:** Advances in aging studies revealed the need to extend the old-age health and functionality data from the *Drosophila-C.elegans-*yeast triad of model organisms to include those that combine the advantages of non-vertebrate models with stronger similarities to mammalian aging. This paper presents a previously unreported phenomenon: reproductive “rejuvenation” in *Daphnia* – a classic and a re-emerging choice model for longevity studies. These postsenescence reproductive episodes result in fecundity similar to that of young individuals and are accompanied by transcriptional shifts, often reverting age-related changes to youthful levels. Transcripts that show such reversal point to restoration of aggregate removal and of mitochondrial function. We believe that this finding opens a novel window of research towards mechanistic understanding of healthy aging.

## Introduction

Scheduling the reproductive effort along the lifespan is one of the key life-history decisions organisms make to maximize their lifetime fitness. While generally selection supports higher investment in early reproduction, as earlier produced offspring have a higher reproductive value (Fisher 1930; Caswell 1982), in certain situations higher late-life reproduction output is observed. Typically, late-life outbursts of reproduction are described as terminal investment: a switch of resource allocation to reproduction in the face of anticipated high mortality (Clutton-Brock, 1984; Forslund & Pärt, 1995; Isaac & Johnson, 2005; Duffield et al. 2017). Essentially this strategy amounts to a late-life switch from iteroparous to semelparous life history. The expectation of increased mortality may be either due to general properties of actuarial aging (e.g., Gompertz law [Olshansky & Carnes 1997; Kirkwood 2015]), or due to environmental changes, for example seasonal, or may be a self-fulfilling prophecy, as redirecting resources for a bout of reproduction may take resources away from maintenance and repair (Duffield et al. 2017). An alternative explanation of late-life reproduction bouts is the exit from a reproductive diapause or otherwise temporary reproductive restriction caused by factors like, again, seasonality, current feeding conditions or time necessary to accumulate resources for successful reproduction (McNamara & Houston 1996). Such periods of reproductive pauses are distinct from senescence and in fact may exist in order to postpone senescence by limiting investment into reproduction to increase survival. Perhaps the most striking example of such rejuvenation is the exit of *Caenorhabditis elegans* larvae from the long-lived, non-reproducing dauer stage (Houthoofd et. al. 2002).

Either way it is of interest for aging research to find out what prior life-history events correlate with late-life reproductive “rejuvenation” and what gene expression changes occur during such a switch, given that as one of aging research and gerontology central goals is to investigate biological foundations of healthy and productive late-age life. Current knowledge about mechanisms of reproductive rejuvenation is largely built around three interrelated hypotheses: that reproductive rejuvenation occurs through the reversal of mitochondrial dysfunction in the germline (Labarta et al. 2019; Chiang et al. 2020), or through attenuation of germ-line apoptosis (Zhao et al. 2017), or through removal of oxidatively damaged lipid or protein aggregates (Terman 2001) in both somatic and germline cells. These likely mechanisms of rejuvenation provide an *a priory* list of functions expected to be regulated to allow rejuvenation: mitochondria function and integrity, aggregate degradation mechanisms, including lysosome acidification (Bohnert & Kenyon 2017), and apoptosis regulation such as the p53 pathway.

In this study we focus on the regularly occurring cases of late-life restoration of reproductive function in *Daphnia -* a freshwater planktonic crustacean that is a classic model organism in aging and longevity research (Ingle 1933), which is enjoying a revival due to contemporary genomic tools (Miner et al. 2012), generating a significant new data on aging (Schumpert et al. 2015; Constantinou et al. 2019; Cai et al. 2020; Hearn et a; 2021; Nguyen et al. 2021; Anderson et al. 2022; Cho et al 2022). *Daphnia* are particularly suitable for cohort longevity and reproductive senescence studies because their reproduction mode, the cyclic parthenogenesis, allows to create cohorts of genetically identical and yet fully outbred individuals, thus providing an advantage over the use of inbred lined customary in *Drosophila* or *C. elegans* longevity studies. Additionally, *Daphnia* transparent bodies allow straightforward measurements of fecundity and lipid content, as well as of the in-vivo use of various tissue-specific fluorescence of dyes.

Our previous studies on aging hallmarks in *Daphnia* indicate that mitochondrial dysfunction is an unlikely cause of aging (and therefore its reversal an unlikely cause of reproductive rejuvenation), as little if any reduction of respiratory activity of mitochondrial membrane potential occurs at old age (Anderson et al. 2022). On the other hand, accumulation of oxidized lipids and misfolded proteins in the forms of lipofucsins and amyloids has been shown to increase with age (Lowman & Yampolsky 2023). Little is known about the role of apoptosis in *Daphnia* aging.

One pattern of *Daphnia* life history that became apparent from a number of longevity studies in this organism is the lack of genetic trade-off between survival and fecundity (Dudycha & Tessier 1999; Coggins et al. 2021). Rather, differences in longevity and fecundity among genotypes are readily explained by differences in genetic load (Lohr et al. 2014; Coggins et al. 2021), while the differences observed within a single genotype reared under different resource availability often emerge through higher, not lower investment into reproduction, relative to growth under limited resources (Lynch and Ennis 1983; Nguyen et al. 2021). Biochemical interventions known to extend lifespan, such as exposure to beta-hydroxybutirate or nicotineamide mononucleotide, also tend to positively affect fecundity rather than longevity (Pearson et al., in preparation). Thus, it appears that *Daphnia* has been under selection to invest into reproduction whenever such opportunity presents itself, at the expense of aging-related investments and life-history consequences.

To focus on possible mechanisms of old-age reproductive rejuvenation in *Daphnia* we first demonstrate that the previously sporadically observed cases of old age restoration of reproductive function (which we term post-senescence reproductive events, PSRE) represent an ubiquitous phenomenon in the long-lived species *D. magna.* We also show that while the reproductive rebound events do occur on the background of exponentially increasing late-life mortality, they neither cause nor anticipate elevated mortality or shorter lifespan in the individual undergoing such reversal of reproductive senescence. We then report the results of an RNAseq differential expression (DE) experiment comparing transcriptomes of young reproducing, senescent post-reproductive, and PSRE females. We discuss functionality of genes with significant differential expression, both in the light of the a priori predicted reproductive senescence reversal pathways discussed above, and focusing on enriched gene ontologies not a priori predicted to play a role in rejuvenation.

## Materials and Methods

Four geographically distant *Daphnia* clones were obtained from Basel University *Daphnia* stock collection and cultivated in the laboratory for at least 5 years. Clones’ identity and geographic origin are listed in Supplementary Table 1. These clones were chosen to represent (2 of each) the long-lived clones from permanent water bodies and short-lived clones from intermittent water bodies (Dudycha & Tessier 1999). Briefly, clones were maintained at the density of 1 adult per 20 mL of water, fed with *Tetradesmus obliquus* (formerly *Scenedesmus acutus*) culture added daily at 100,000 cells/mL. Further details on clones’ provenance and maintenance in Anderson et al. 2022.

Experimental animals were collected as neonates (less than 24 h old, all females) from 15-25 days old mothers and placed individually in 35 mL shell vials containing 20 mL of water. The vials were maintained daily and age and body size at maturity were recorded. Subsequently, neonates were counted and removed, and water was changed at the end of each ovary cycle (4 days at 20^°^C, 2-3 days at 24^°^C). Each vial was maintained until the death of the individual. Two separate experiments were conducted, referred to as Experiments 1 and 2 thereafter.

Experiment 1 consisted of 94 individuals kept in COMBO medium (Kilham et al. 1998) at 20 ^°^C. Experiment 2 included three treatments: COMBO medium at 20 ^°^C as in Experiment 1, ADaM medium (Klüttgen et al. 1994) at 20 ^°^C, and ADaM medium at 24 ^°^C, with 89, 91, and 88 individuals in each of these 3 cohorts. The two media are similar, both imitating ionic composition of natural pond water, but ADaM contains significantly higher concentration of Ca^2+^ and Mg^2+^ ions, modeling water with higher hardness.

Reproductive “rejuvenation” (post-senescence reproductive events, PSRE) was defined as a production of a clutch of at least 6 live neonates such that the number of neonates produced was at least 3 standard deviations above the mean number of neonates produced in the previous 5 ovary cycles (including any cycles in which no neonates were produced) by a female over 60 days of age. Subsequent large clutches meeting the same criterion were considered a continuation of the same PSRE. In a few cases two or even three PSRE events occurred in the same individual, separated by several low-or no reproduction cycles; these were counted as separate events. Differences of life-time survival among clones and treatments as well as in post-PSRE survival between individuals showing a PSRE event and matching non-rejuvenated individuals were analyzed by means of Kaplan-Meyer and proportional hazards methods implemented in JMP statistical platform (ver. 17; SAS Institute 2018).

Starting with age of 30 days random samples of neonates born to mothers in the ADaM medium, 24 ^°^C treatment were taken for neonate size and lipid content measurement. Neonates were sampled <24 hours after birth and stained with Nile Red (1 µg/mL) for 1 hour. After staining neonates were photographed individually using a Leica DM3000 microscope with a 10× objective (0.22 aperture) equipped with a Leica DFc450C color camera, with a 488 nm excitation/broadband (>515 nm) emission filter (for Nile Red fluorescence) and in bright field (for body length measurement). Images were analyzed using ImageJ (Rasband et al. 2018) with the portion of intensity emitted from areas with unweighted intensity over 150 recorded as the measure of lipid content (see example image in Results section). An additional cohort was started at day 30 of the experiment to allow a common garden comparison of neonates born to younger and older individuals.

Four 130 - 175 days old GB-EL75-69 females from the ADaM medium, 20 ^°^C treatment that have just produced a clutch meeting the PSRE criterion described above were removed from the experiment and flash-frozen in liquid nitrogen for the RNAseq experiment. Four randomly chosen reproductively senescent females from the same clone and treatment were selected as the senescent controls. These females were carrying no clutch or, in one case, carrying a clutch of just 2 eggs. These two samples will be hereon referred to as Old, PSRE and Old, non-PSRE. For the Young sample four randomly chosen individuals from the same clone from a separate cohort of the age on 20 days were treated in the same manner. In all cases the individuals were sampled within 24 hours of producing a clutch or from the last molting if no clutch was produced. The eggs were removed from the brood chamber prior to freezing and frozen separately.

Whole organism RNAs were extracted from these individuals using Qiagen RNeasy Plus Mini kit. RNA library preparation performed using NEBNext Ultra II Directional RNA Library prep Kit (NEB, Lynn, MA) following the manufacturer’s protocols. The libraries were sequenced with Illumina Novoseq 6000, S4 flow cell, PE100. The quality check of Indexed sequences was performed by Fastqc, and indexed sequences were trimmed using an adaptor sequence by TrimGalore-0.4.5.

Reads were mapped to *D. magna* Xinb3 reference transcriptome (BioProject ID: PRJNA624896; D. Ebert and P. Fields, personal communication; Fields et al., in preparation) using STAR (Dobin et al. 2013) and genes with differential expression (DE) either between young and old *Daphnia* or between PSRE- and non-PSRE old *Daphnia* were identified using DEseq2 (Love et al. 2014). To analyze Gene Ontology (GO) enrichment among genes with a significant DE by the list of such genes, in each comparison separately, was matching to the list of GOs and pathways identified in reference *D. magna* transcriptome obtained by PANTHER (Ver. 16; Mi et al. 2019; Thomas et al. 2022). Counts of genes representing each GO or pathway in a list consisting of unique combinations of genes and GOs was then tested for being heterogeneous (enriched or depleted) relative to the numbers of other genes in a given gene list and the numbers of other GOs in the reference using Fisher’s Exact Test (FET). The four counts used in the FET for each GO category were the number of genes in this GO category with DE; the number of genes in this category without DE, the number of genes in all other GO categories in the reference set showing DE, and the number of genes in all other GO categories in the reference set without DE. False Discovery Rate adjustment for multiple testing was applied and only the innermost nest GO category showing a significant enrichment or depletion is reported.

## Results

### Occurrence of post-senescence reproductive events

Overall lifespan and age-specific survival differed significantly between the two experiments, two temperatures within experiment 2, and among clones in experiment 2 (Supplementary Fig. S2, Supplementary Table S3). There was, however, no significant differences between the two media tested in Experiment 2. Despite these overall differences, PSRE’s occurred in nearly all combinations of clones and treatments with frequencies of occurance ranging from 10 to 60% of survivors to (Table 1).

**Table 1.**
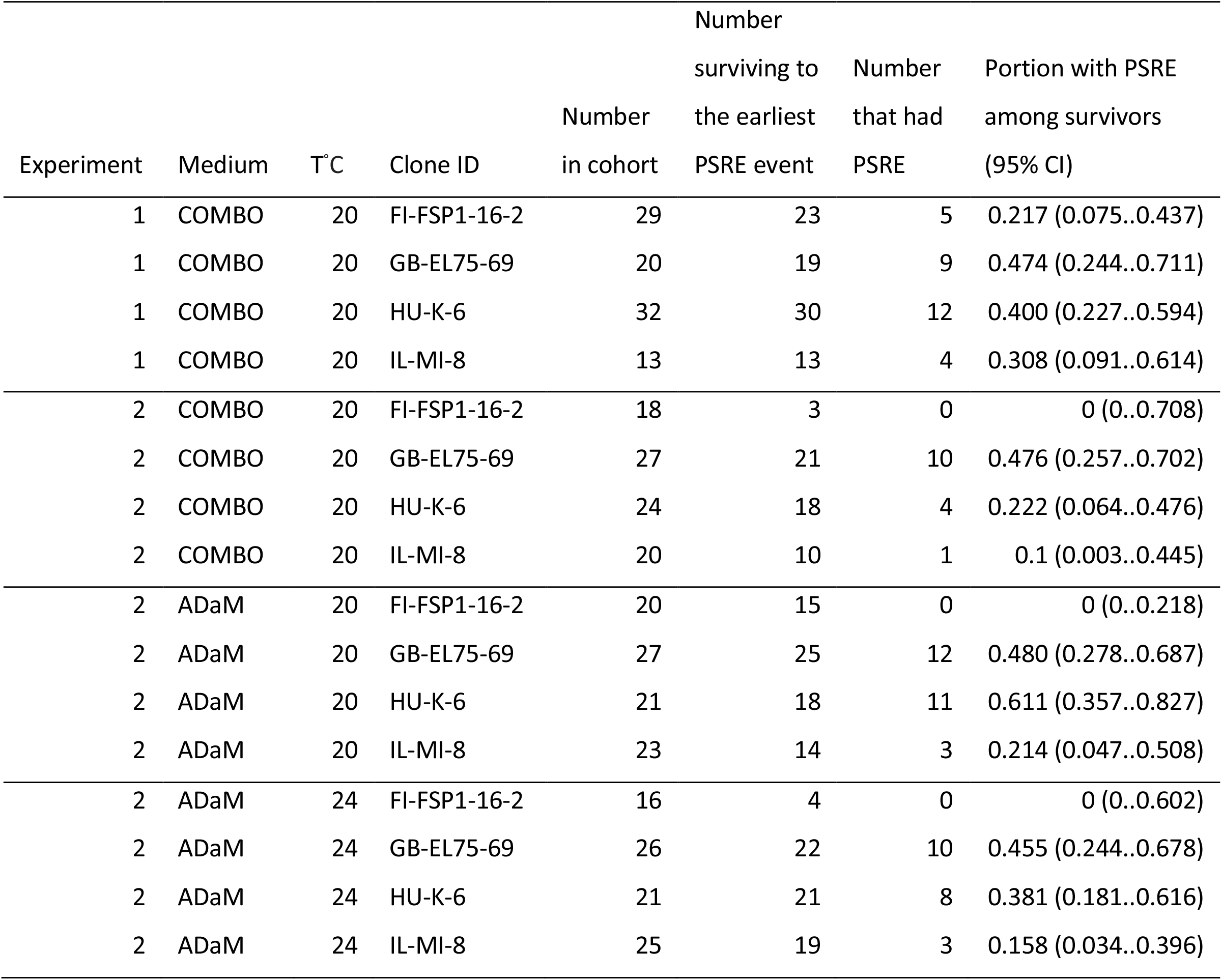
Frequency of post-senescence reproduction events (PSRE) in *Daphnia* from 4 clones in life-history experiments conducted in various conditions. 2-way nominal logistic fit p-value for the difference among conditions (combination of medium and temperature): p<0.01; for the difference among clones: p<0.001; for the interaction: p>0.45. CI: Clopper–Pearson CI.

### Trade-offs with late-life mortality and investments per offspring

While the PSRE events occurred during period of high mortality and preceded further increase of mortality overall (Fig. 1), we did not detect any increase of mortality following a PSRE event specifically in individuals experiencing such events. In fact, life expectancy of such individuals was higher than of non-PSRE individuals of comparable age (Fig. 2, Table 2). Thus, late-age reproduction neither anticipates nor causes elevated mortality. However, trade-offs were observed between late-life number of offspring (clutch size) and body size and lipid content of offspring (Fig. 3). Offspring body length in PSRE individuals was consistently lower than in their non-PSRE counterparts, both significantly decreasing with age (Fig. 3A), although this difference was small in magnitude (approximately 20 μm or 2% of body length). A different pattern of trade-off was observed for lipid content of the neonates (Fig. 3B). The PSRE females were producing neonates with a higher lipid content than their non-PSRE counterparts at ages 75-90 days, but rapidly reduced lipid investment per offspring at ages over 100 days, indicating higher cost of producing PSRE clutches at older age, as indicated by a significantly higher slope of neonate lipid content over mother’s age in PSRE than in non-PSRE mothers (Fig.3B).

**Fig. 1.**
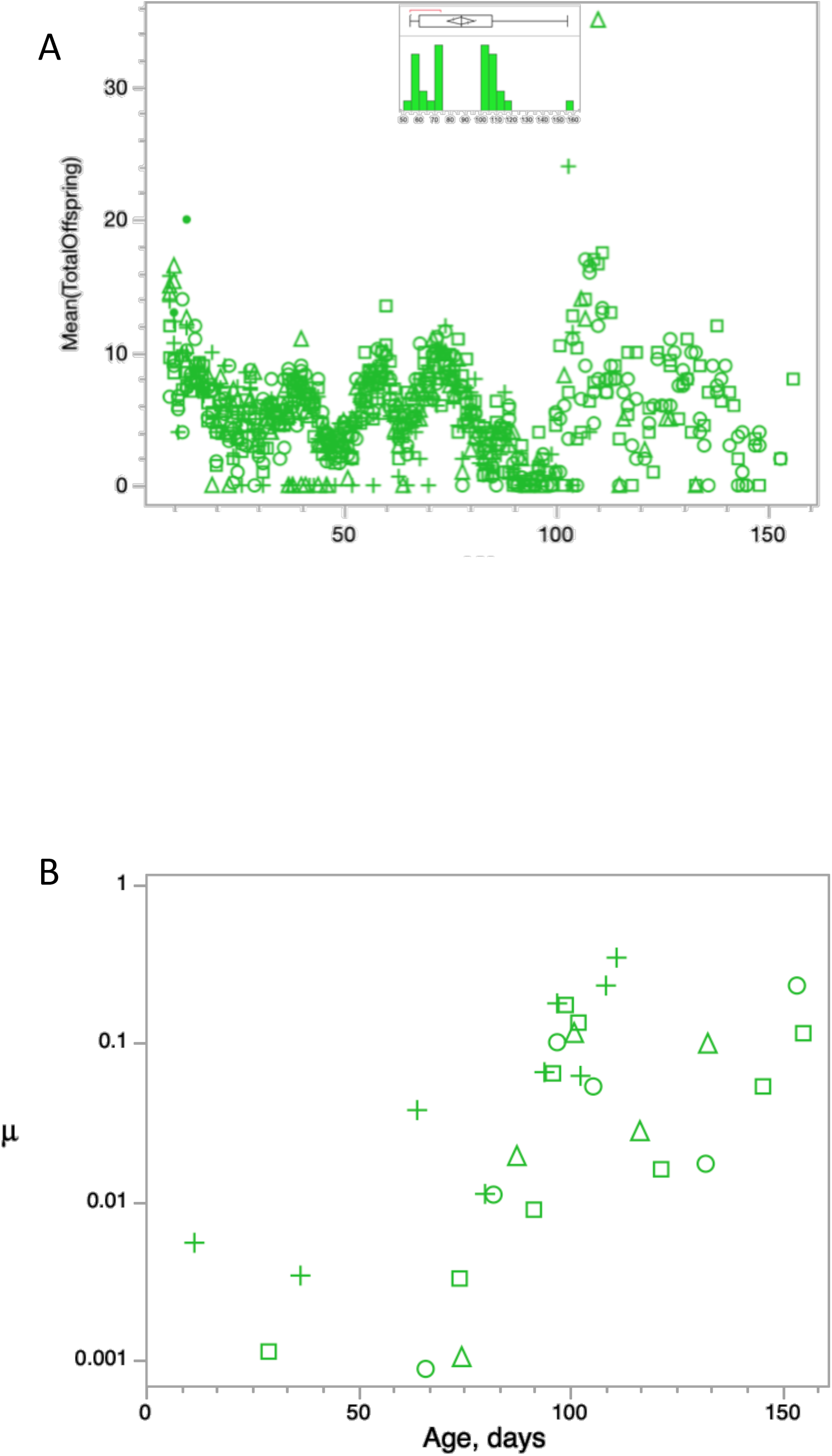
Fecundity (A, C, D, E) and mortality 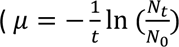, B, G, H, I) in life-table experiments 1 (A,B) and 2 (C-I). Colors correspond to experimental treatments: COMBO medium, 20 °C: green; ADaM medium, 20 °C: blue; ADaM medium, 24 °C: red. Inserts show distribution of PSRE events over age. Symbols correspond to 4 different clones used. See Supplementary Fig. S2 for clone-specific mortality data.

**Fig 2.**
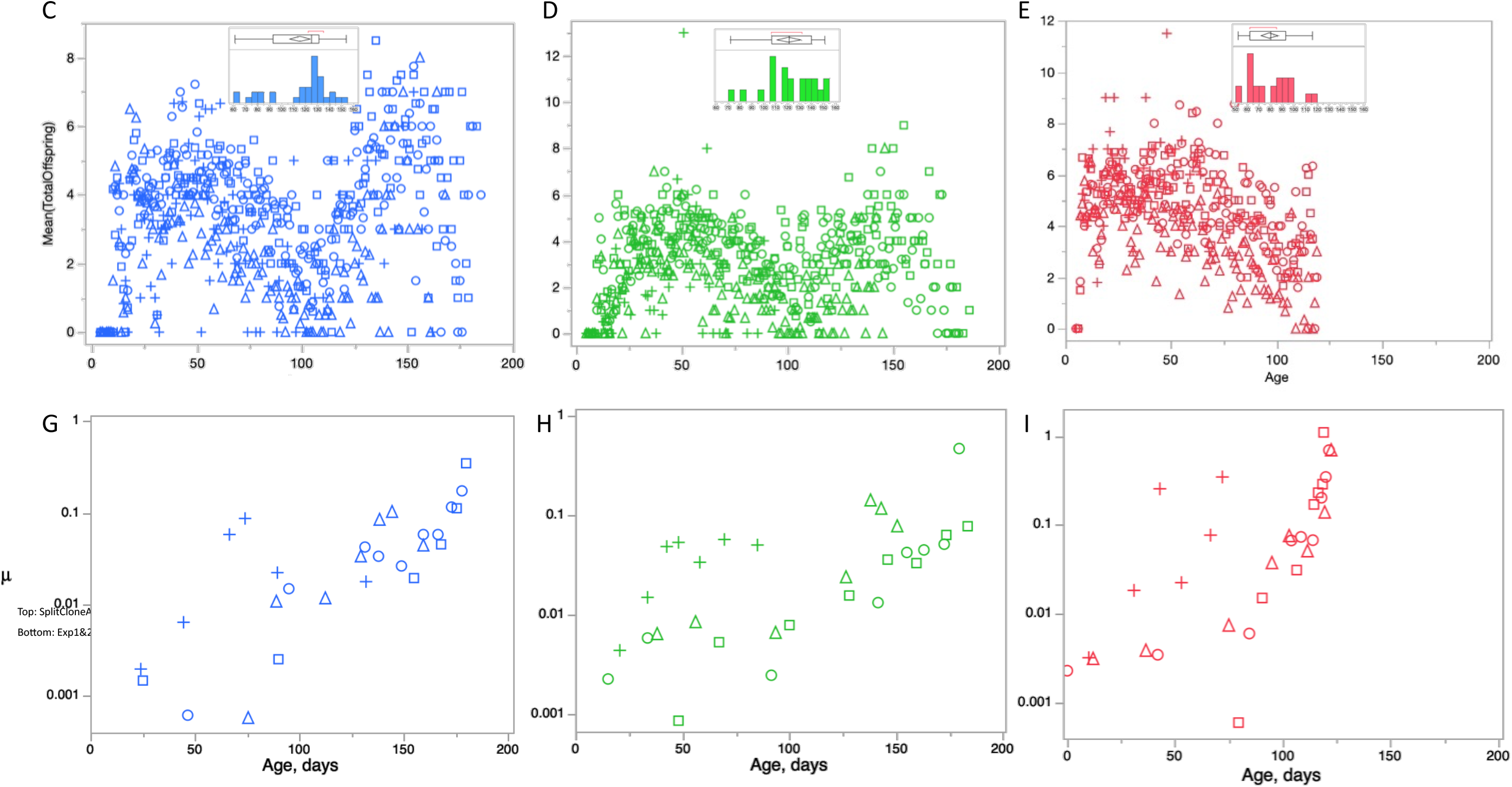
Survival of *Daphnia* past a late-life reproduction bout event (blue) vs. survival of individuals not showing a late-life reproduction bouts, past average age of late-life reproduction bouts in the same clones in the same experiment (red). P-values from Kaplan-Meyer survival analysis: p_L-R_ – Log-rank test; p_W_ - Wilcoxon test. See Table 2 for more detailed analysis.

**Fig 3.**
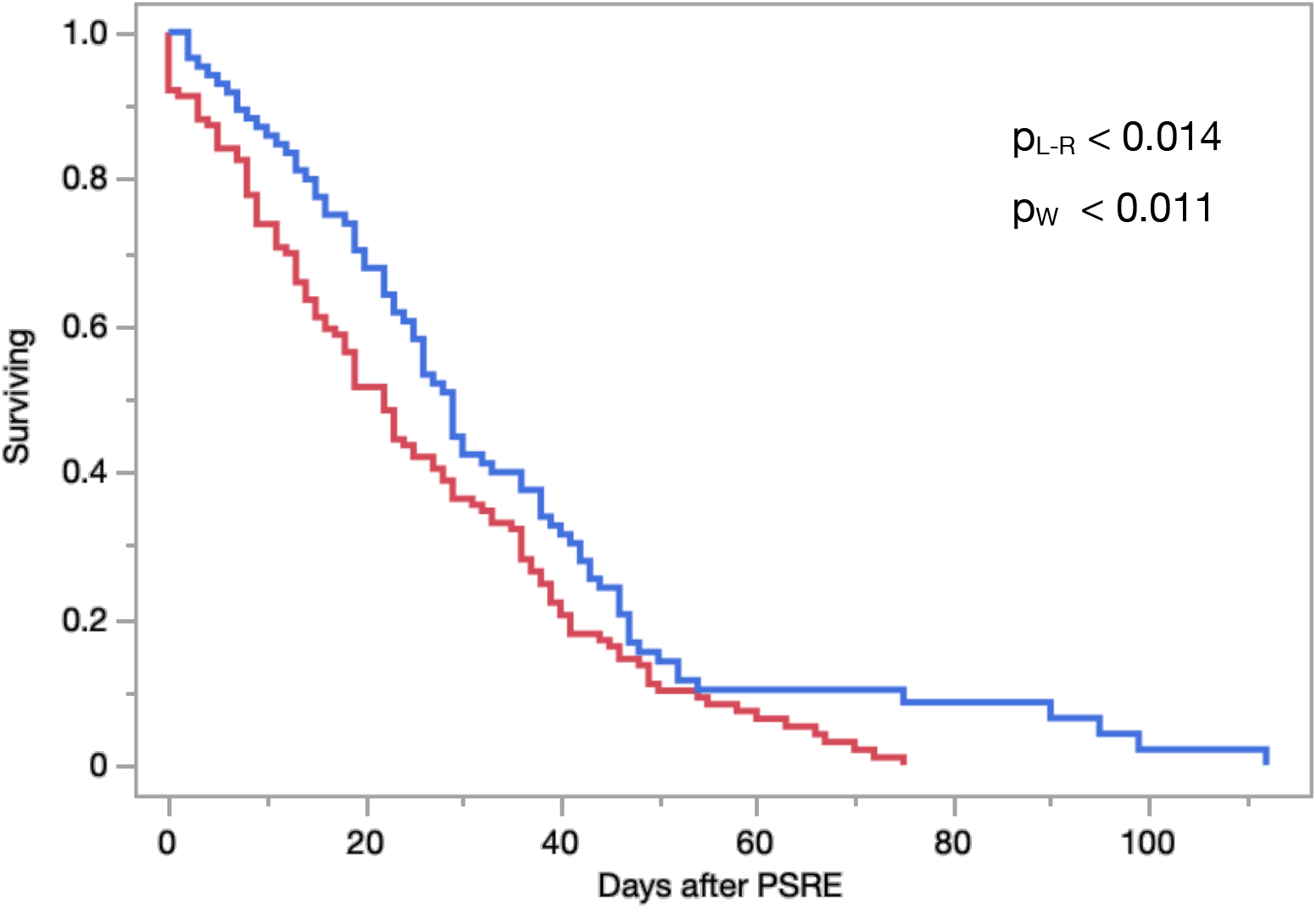
Body length (A) and lipid content (B) in neonates born to Daphnia females of different age who either have or have not experienced a PSRE while producing these neonates (blue and red, respectively). P-values represent heterogeneity of slopes test (A: differences in intercept and slope, B: differences in slope). Neonate parameters were determined only in AdaM 20 °C treatment.

**Table 2.**
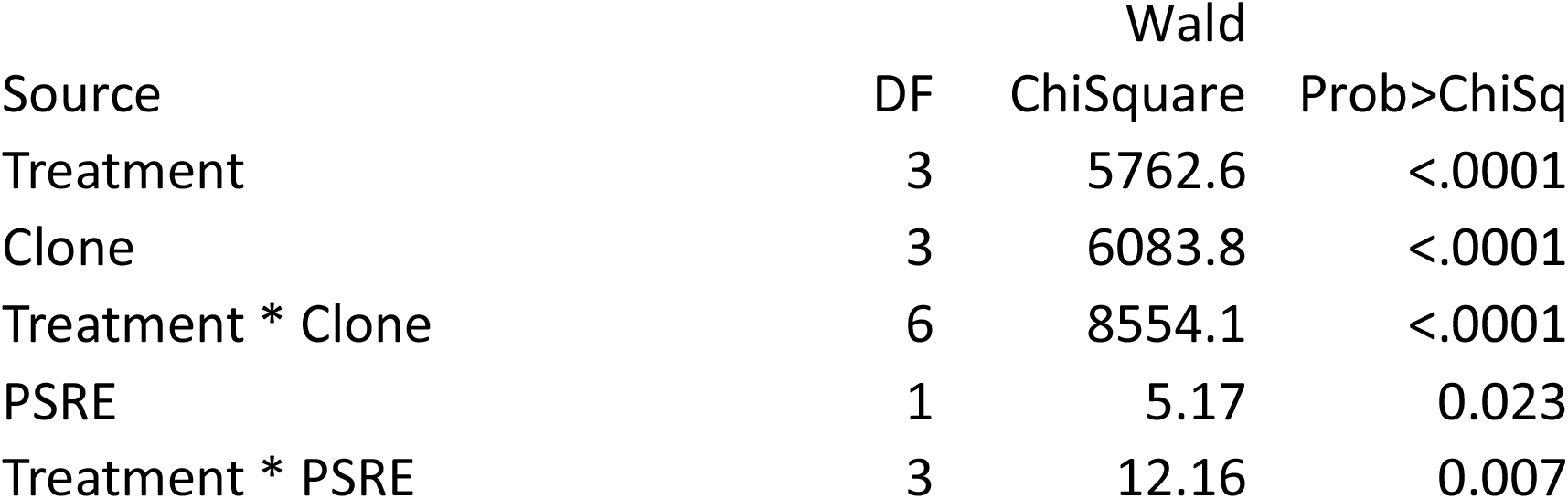
Proportional Hazards analysis of late-life survival of *Daphnia* with and without post-senescence reproductive events (PSRE). Overall survival shown on Fig. 2.

| Source | DF | Wald |  |
| --- | --- | --- | --- |
|  |  | ChiSquare | Prob>ChiSq |
| Treatment | 3 | 5762.6 | <.0001 |
| Clone | 3 | 6083.8 | <.0001 |
| Treatment * Clone | 6 | 8554.1 | <.0001 |
| PSRE | 1 | 5.17 | 0.023 |
| Treatment * PSRE | 3 | 12.16 | 0.007 |

### Correlations with environment and early life events

In order to determine if early-age life history events predetermine late-life propensity to exhibit PSRE event we correlated early body size at maturity and fecundity (mean clutch size) in age classes <10, 10-30, 30-50 and 50-70 days with future PSRE event in Experiment 2 (Fig. 4). In 2 out of the 3 clones that showed PSRE events in Experiment 2 individuals were smaller at maturity than those that survived to the minimal PSRE age, but never experiences a PSRE (Fig. 4A); overall effect was not significant, but Clone-by-body size interaction was (Supplementary Table S3). Clutch sizes in ages 10-70 were not a predictor of future PSRE (Fig. 4B, Supplementary Table S4), except for a border-line significant effect of fecundity in the age class 10-30 days (non-significant after multiple test correction) despite apparently slightly higher fecundity in future PSRE females than their same-age counterparts (Fig. 4B, Supplementary Fig. S2).

**Fig 4.**
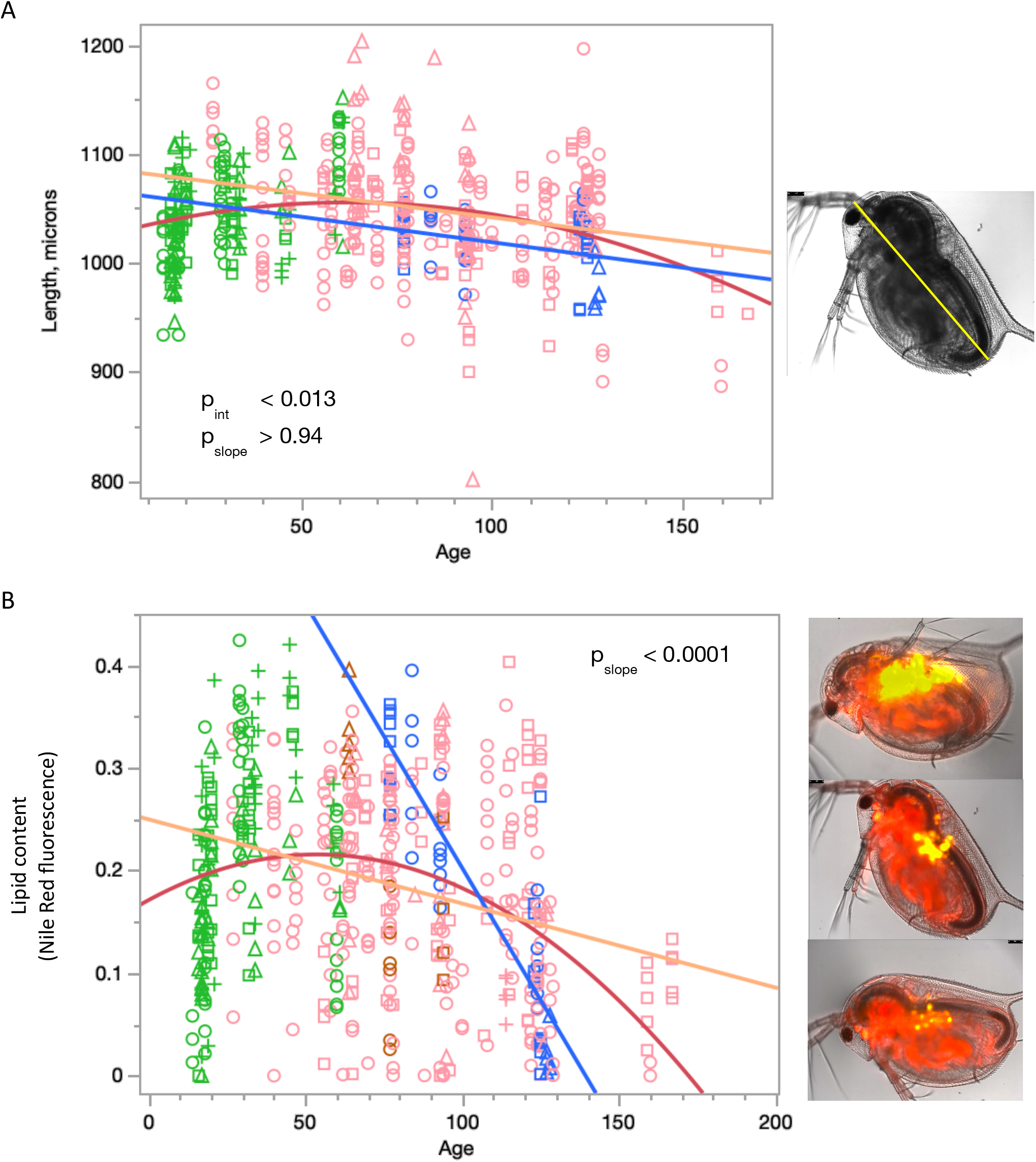
Body length at maturity (A) and early-to-mid life fecundity (clutch size) in Daphnia females who either would (blue) or would not (red) experience a PSRE later in life. Only females who survived to the earliest age of PRSE are included into this analysis. P-values represent the PSRE status effect in a 3-way nominal logistic regression with treatments, clones and either body size or early fecundity on future PSRE (Supplementary Tables S2 and S3). Non-PSRE clone FI included for comparison. See Supplementary Fig. S2 for combined data across both experiments.

### RNAseq results

Transcriptional profiles differed between young, old non-PSRE, and old PSRE individuals. Principal component analysis revealed a good separation between the three groups along the 1^st^ PC that explained 50.7% of the variance, with the old PSRE group positioned between the young and old non-reproducing individual samples (Fig 5). PC 2 (14.7%) separated, with one sample exception, the reproducing individuals from the non-reproducing ones, indicating either overcompensation in PSRE individuals or otherwise PSRE-specific transcription. Much of the differential expression manifested through overexpression of numerous transcripts in old, relative to young individuals and in non-PSRE vs PSRE individuals (Supplementary Fig. S3).

**Fig. 5.**
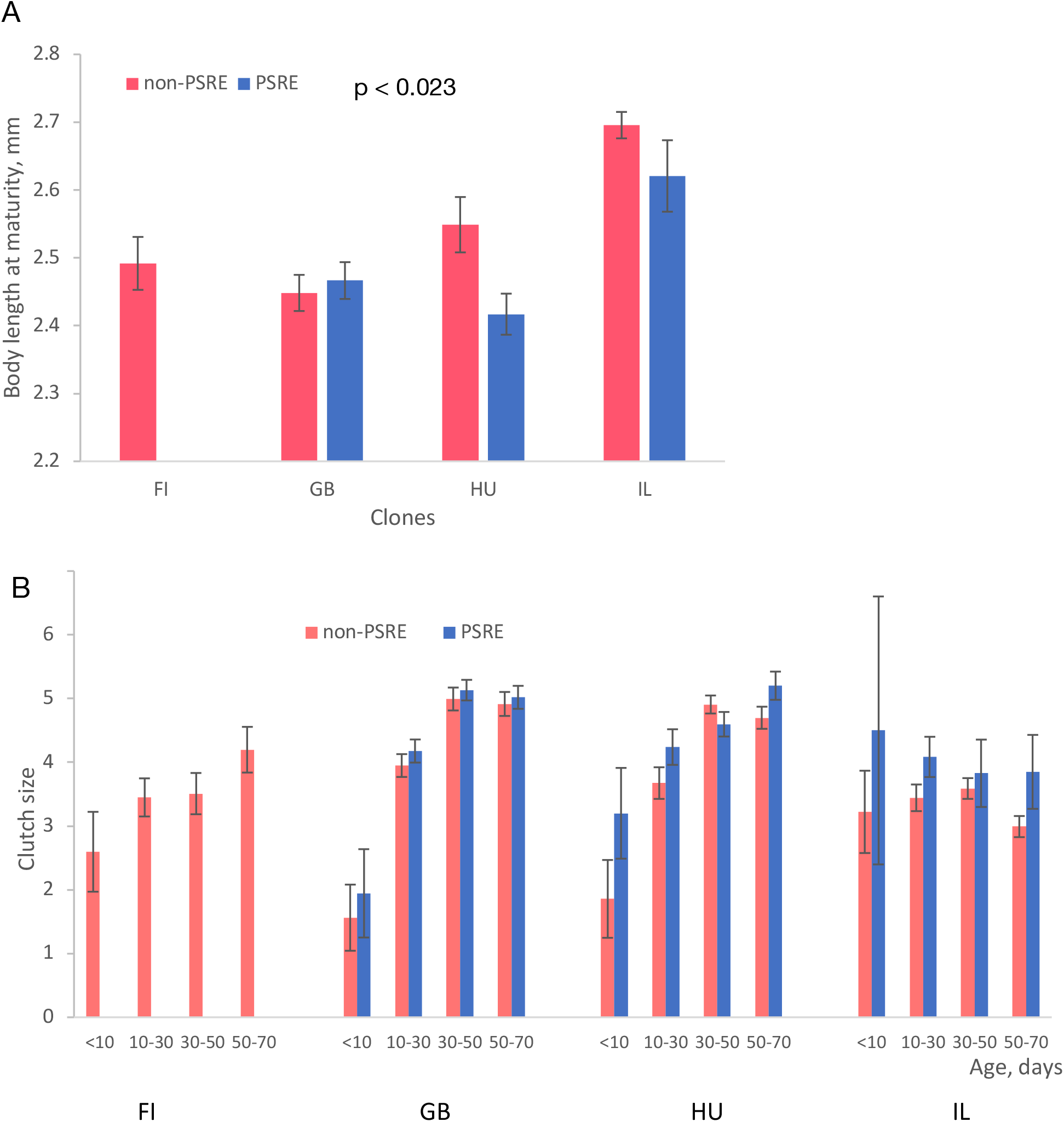
Principal components analysis of young (green), old non-PSRE (orange), and old, PSRE (purple) *Daphnia* based on 1658 transcripts with a significant DE (p_adj_<0.05; |logFC|>1) between age classes.

GO enrichment analysis results are shown in Supplementary Table S4. Many of the GO or pathway terms listed were enriched among differentially expressed due to enrichment of a particular gene family, rather than unrelated genes with different functionality within the same GO term. Expression profiles of several such gene families are shown on Figs 6 and 7. For the genes up-regulated in old, nonreproducing *Daphnia* (Fig. 6) these gene families included guanidine monophosphate synthases (GMPSs), guanylate cyclases (GCs), as well as two putative transposon-related gene families (see below). In contrast, genes showing down-regulation in old *Daphnia* relative to the young ones were enriched in GO terms related to RNA processing, ribosome structure, and protein synthesis, most notably in ribosomal proteins (Supplementary Table S4).

**Fig. 6.**
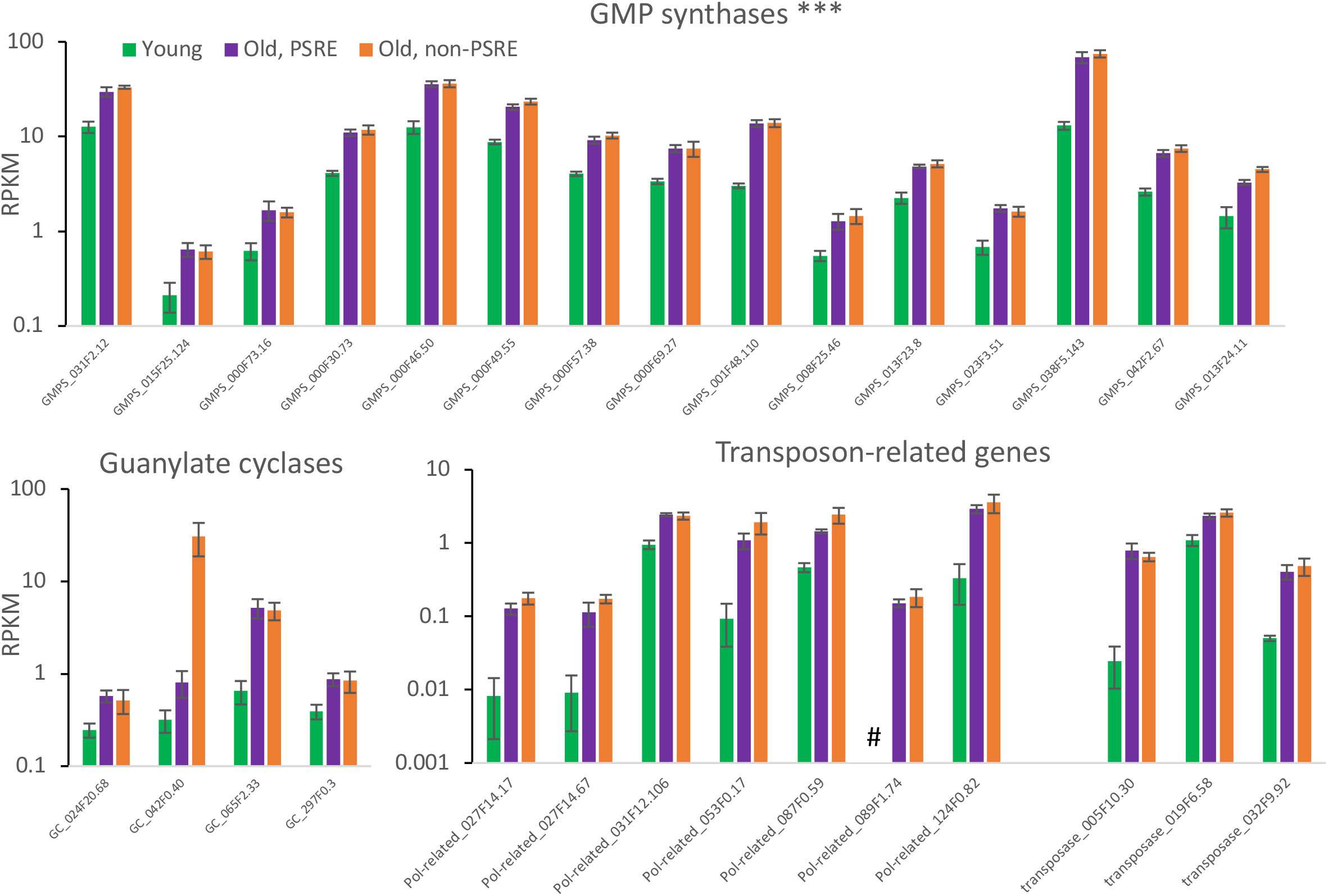
Expression level (RPKM, log scale) of select gene families or GOs of interest in young (green), old PSRE (purple), and old non-PSRE (orange) *Daphnia* showing DE with at least P_adj_<0.01 and at least |log_2_(FC)|>1 between young and old *Daphnia*, but not between PSRE and non-PSRE categories. Bars represent SE among the four samples and do not correspond to P_adj_ values estimated by DESeq2. GO significantly enriched among genes with DE (*** p<0.0001; Supplementary Table S4). #: RPKM = 0 (no reads observed).

**Fig. 7.**
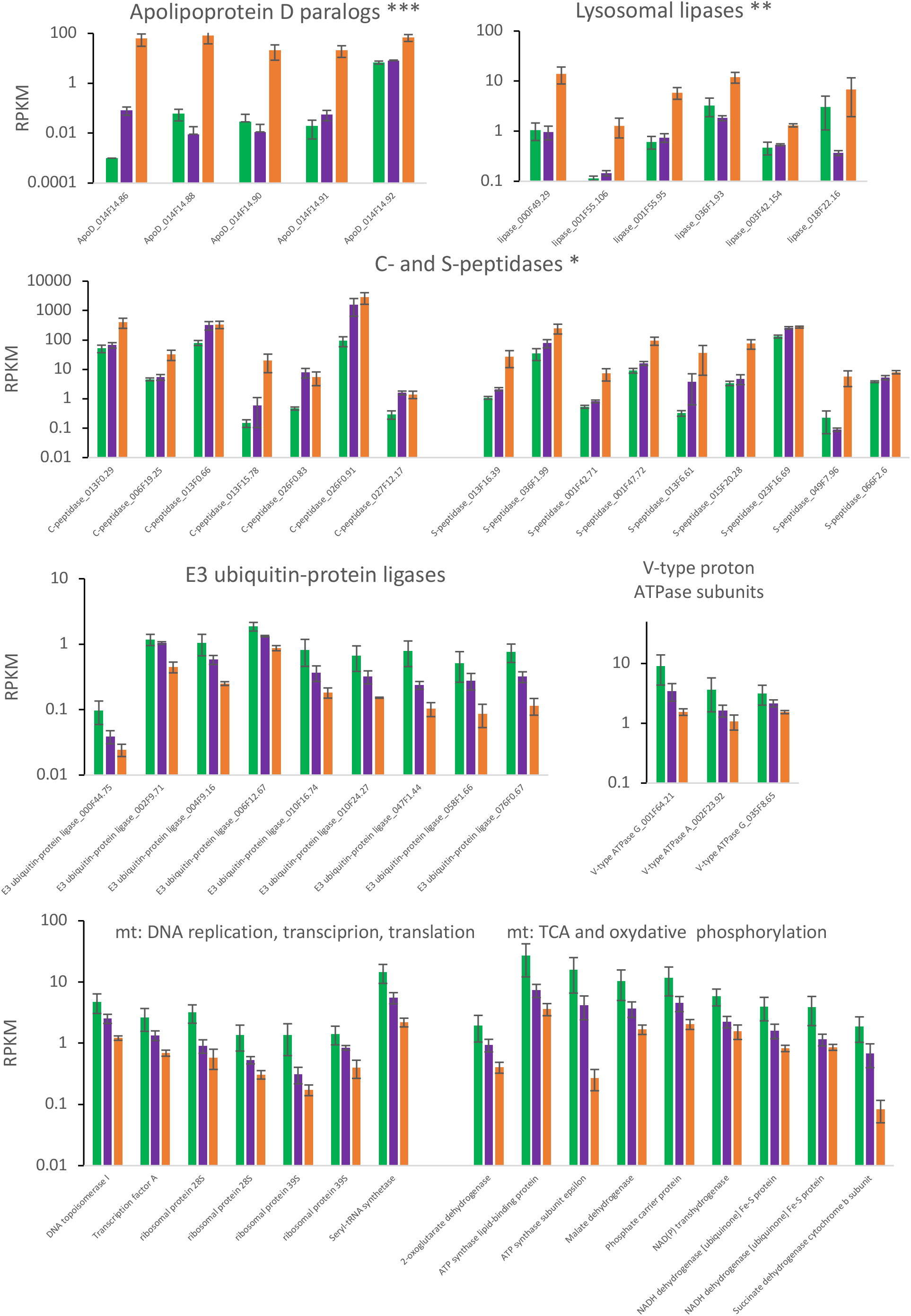
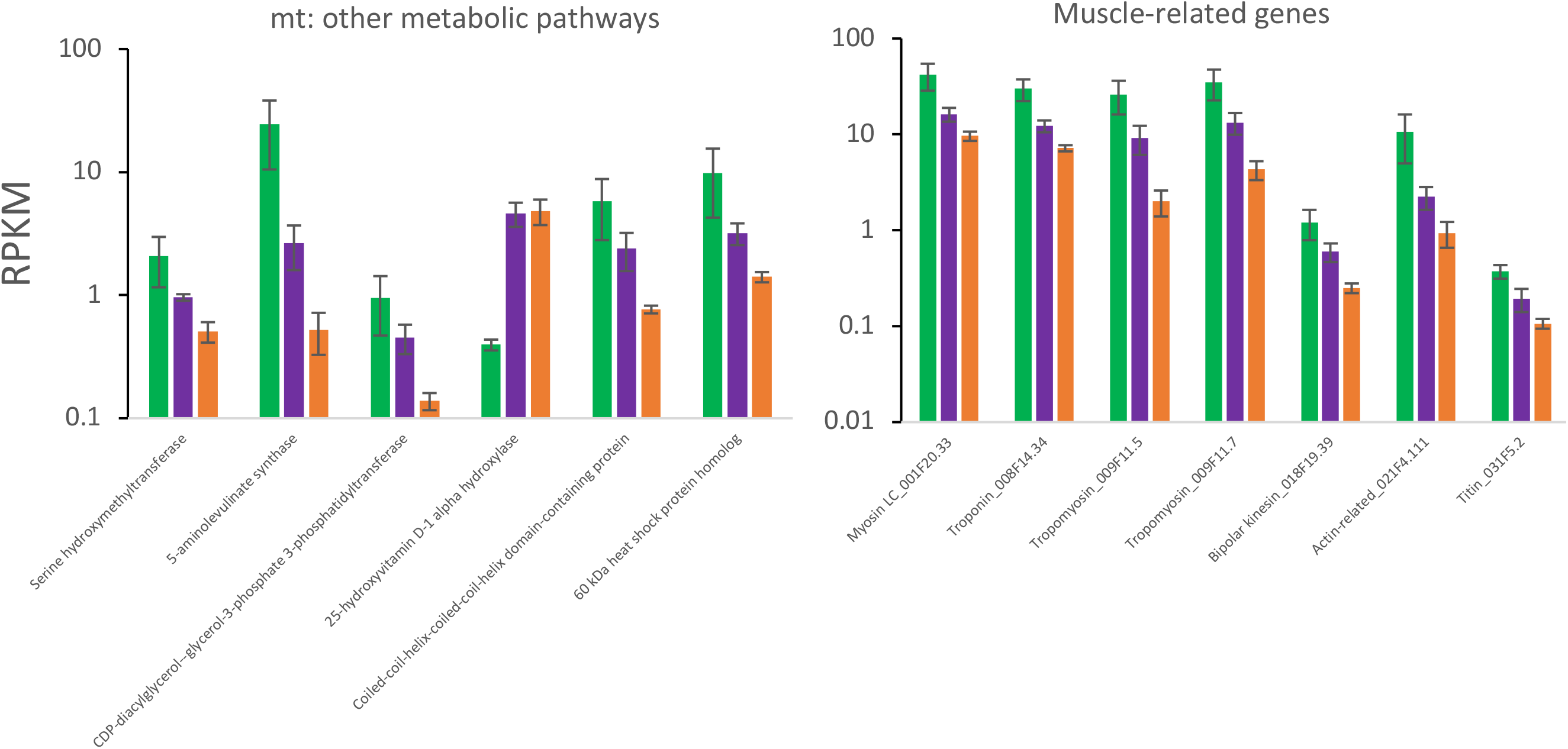
Expression level (RPKM, log scale) of select gene families or GOs of interest in young (green), old PSRE (purple), and old non-PSRE (orange) *Daphnia* showing DE with at least P_adj_<0.01 and at least |log_2_(FC)|>1 between young and old non-PSRE *Daphnia*, with PSRE individuals showing full or partial reversal of transcription to youthful levels. Bars represent SE among the four samples and do not correspond to P_adj_ values estimated by DESeq2. GO significantly enriched among genes with DE (* p<0.01; ** p<0.001; *** p<0.0001; Supplementary Table S4).

GOs and pathways enriched among genes showing DE between PSRE and nonPSRE old *Daphnia* are shown on Fig. 7. These include apolipoproteins D (apoD), two families of endopeptidases and a family of lysosomic lipases. Notably, in PSRE old *Dapnnia,* all of the apoD’s, and several of lipases and endopeptidases show full or partial reversal back to low level of expression characteristic to young *Daphnia.* The same pattern is observed in several gene families and functional groups not detected as enriched by the GO enrichment analysis (see below).

Differentially expressed members of each gene family included both tandemly or segmentally duplicated and unlinked paralogs. However, all old age-specific apoD’s are tightly linked, mapping DM3 assembly scaffold 0014F, scaffold positions 14.86-14.92. The the greatly expanded GMPS gene family includes a series of paralogs mapping to scaffold 0000F, with scaffold positions between 30.73 and 73.16 and a pair of linked paralogs located in the scaffold 0013F, positions 23.8 and 24.11. Old age-specific lysosomal lipases include a pair of closely linked paralogs in scaffold 001F positions 55.95 and 55.106 and several gene family members located elsewhere.

It is important to notice that the lists of differentially expressed genes are rich in genes with no GO or pathway terms that could be assigned to them, with many of such genes being *Daphnia-*specific. Unexpectedly, the lists of up- and down-regulated genes (DESeq2 adjusted P<0.01) differed with respect to the abundance of such genes. For the young vs. old, nonreproducing *Daphnia* comparisons the genes up-regulated in the old *Daphnia* were significantly enriched with uncharacterized genes (expected: 315.2, observed: 464, 2-tailed uncorrected Fisher Exact Test P-value <0.0001), while in the list of genes down-regulated in the old *Daphnia* such genes were depleted (expected: 232.8, observed: 91, P <0.0001). In the old, reproducing (PSRE) vs. old, non-reproducing *Daphnia* the list of genes up-regulated in the old, non-reproducing *Daphnia* was slightly enriched in the uncharacterized genes (expected: 75.0, observed: 93, P < 0.008), while the much smaller list of the down-regulated genes was neither depleted nor significantly enriched (expected: 5.8, observed: 8, P>0.17).

All genes with significant DE between young, old non-reproducing, and old reproducing *Daphnia* not already detected as parts of enriched pathways of GO by the above analysis are listed in Supplementary tables S5-S7. They appear to represent a variety of metabolic and regulatory pathways with little to no correlation to a priori expectations. One exception is a noticeable up-regulation of putative transposon-related genes in old, non-reproducing *Daphnia.* Both retro-transposons and DNA transposons are represented (6 paralogs of retrotransposon-related homologs of *pol* protein and 3 transposases, respectively, Supplementary Table S5, Figure 7). Another group of genes with predictable age effect not picked up by the GO analysis are those encoding muscle- and motility-related proteins, several of which are down-regulated in old non-reproducing flies, including troponin, tropomyosin alpha-1, bipolar kinesin, titin and myosin light chain (Supplementary Table S6; Figure 7). Furthermore, consistent with the apoptosis- and the aggregates removal predictions about rejuvenation, the same patterns of at least partial reversal of age-related transcriptional changes in the PSRE *Daphnia* is observed in several E3 ubiquitin-protein ligases and several subunits of V-type proton ATPases. Notably, another large group of proteins showing the pattern of transcription level reversal in the PSRE *Daphnia* are proteins with mitochondria-related functionality (Fig. 7). These include proteins directly involved in TCA and/or membrane phosphorylation (including succinate dehydrogenase cytochrome B and subunits of NADH dehydrogenase and ATP synthase) and proteins involved in mitochondrial DNA replication, transcription, and translation, as well a variety of proteins accomplishing other mitochondrial functions (Fig. 7), including, among others, the 5-aminolevulinate synthase catalyzing the first step of heme synthesis and one of mitochondrial heat shock proteins. All but one transcripts in this category show reduced transcription with age ful partially restored in the PSRE individuals.

Finally, several groups of paralogs showed an apparent subfunctionalization by age, with some paralogs being up- and others down-regulated in old, non-reproducing flies (Supplementary Table S7, Fig. 8). Notably, several of these paralogs, including mitochondrial cytochromes P450, chorion peroxidases and zinc finger proteins also show DE between reproducing and non-reproducing old *Daphnia,* indicating that age-specialization of paralogs may be not only related to absolute age, but also to retention of old-age reproductive ability.

**Fig. 8.**
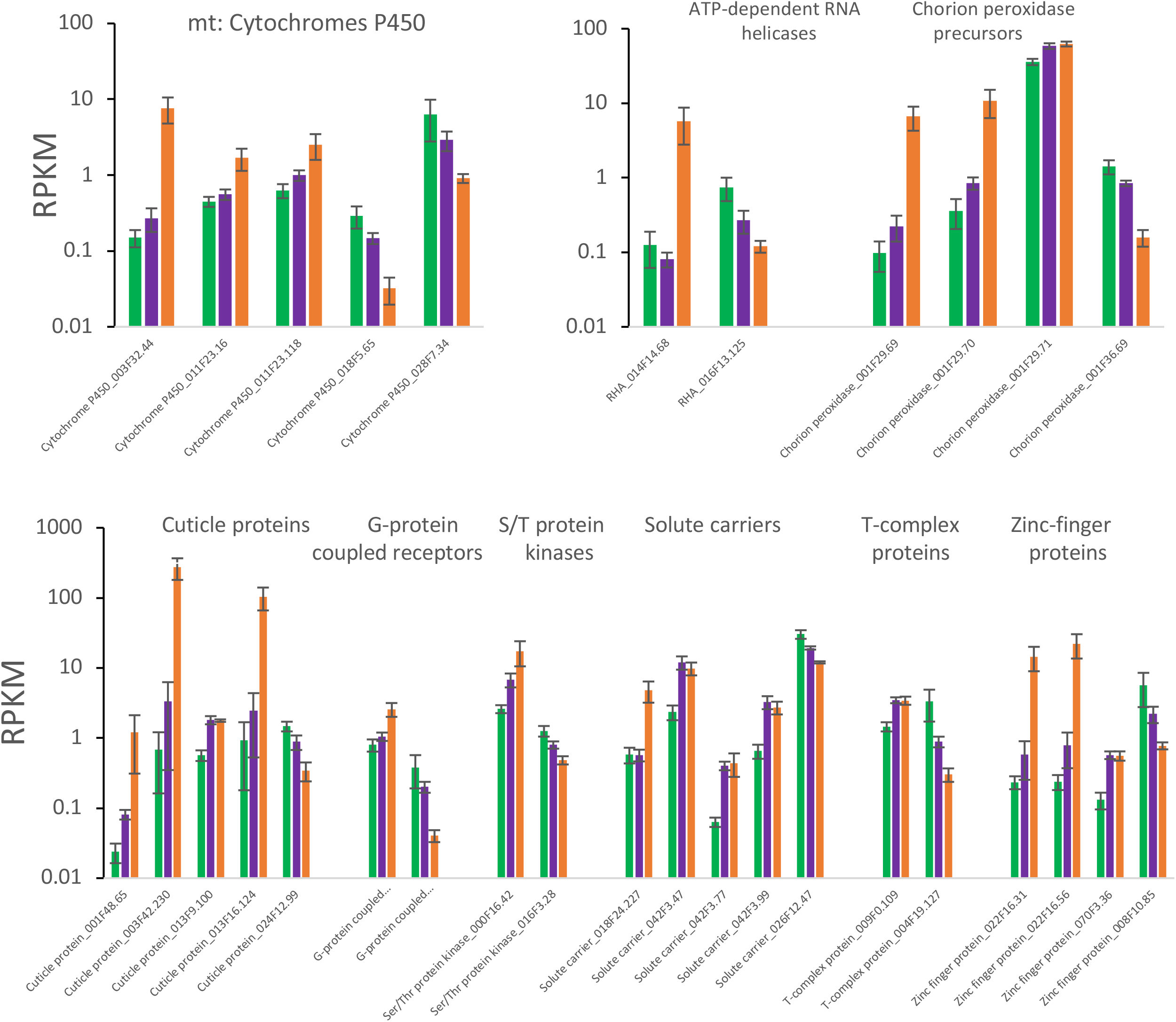
Expression level (RPKM, log scale) of selected gene families with opposite direction of age-related DE in different paralogs. Genes with DE with at least P_adj_<0.001 and at least |log_2_(FC)|>1 shown. Bars represent SE among the four samples and do not correspond to P_adj_ values estimated by DESeq2.

## Discussion

We have observed nearly ubiquitous cases of renewal of asexual reproduction in a cyclic parthenogen *Daphnia* after complete or nearly complete reproductive senescence. Most of these cases occur at the age of 100-150 days, which is 70-80% of maximal lifespan observed in our experiments and the age to which about 20% of a typical cohort survives. We did not observe any evidence that such renewal of reproduction represents the terminal investment strategy, as individuals with a post-senescence reproduction event show a higher, not lower remaining lifespan. This such events neither anticipate nor causes imminent mortality. We did observe a trade-off with offspring size, however, as the offspring of the oldest mothers producing unusually large clutches are smaller than their small-clutch counterparts. This can be interpreted as shortage of resources necessary for oocyte provisioning older *Daphnia* are experiencing even when their ovaries reactivate. These resources are, however, different from the main nutrient *Daphnia* mothers provision their offspring with – lipids, as there is no difference in lipid content between late-life small and large clutches.

We do not know if the observed phenomenon is an adaptation to seasonal environment.

Perhaps even in the conditions of constant light and temperature, *Daphnia* has an internal clock, allowing interpretation of the advanced age as the signal of having survived a winter and therefore anticipate improved environmental conditions for the offspring. If this is the case, the observed reproductive senescence should be interpreted as reproductive diapause and post-senescence reproductive effort - as the exit from it. It should be noted though that post-senescent reproductive events have been observed not only in the *D.magna* clone GB-EL75-69 extracted from a permanent pond in London, UK, where the clone’s recent evolutionary history very likely included overwintering adults, but also in clones IL-MI-8 and FI-FSP1-16-2 originating from temporary pools in which survival unfavorable seasons as adults is highly unlikely.

We have observed a profound change in transcriptome-wide and GO- and gene family-focused transcriptional changes between young and old *Daphnia,* as well as between old non-reproducing and old, reproducing ones. In many cases the old, reproducing (PSRE) individuals showed a partial or even complete reversal of transcriptional profile back to youthful values. It may be noted, that in the multidimentional analysis the biological replicates representing this condition were more similar to each other than those representing young and old, non-reproducing individual, indicating a coordinated transcriptional state.

On the level of individual genes, gene families, and GOs, the transcripts showing aging-related DE either fit into *a priori* predictions (such as mitochondrial proteins, or proteins involved in apoptosis or lipid and protein aggregate removal), or showed a significant enrichment in the GO enrichment analysis. Functionally, the GOs and gene families implicated here as transcriptional markers of aging and of reproductive reversal represented of mix of functionalities that could and could not be easily expected a priori, based on our knowledge of functional genomics of aging and reproduction in better understood models such as *Drosophila* and *Caenorhabdites.* A graphic summary of the observed changes and their hypothesized consequences for physiology and life history is presented on Fig. 9.

**Fig. 9.**
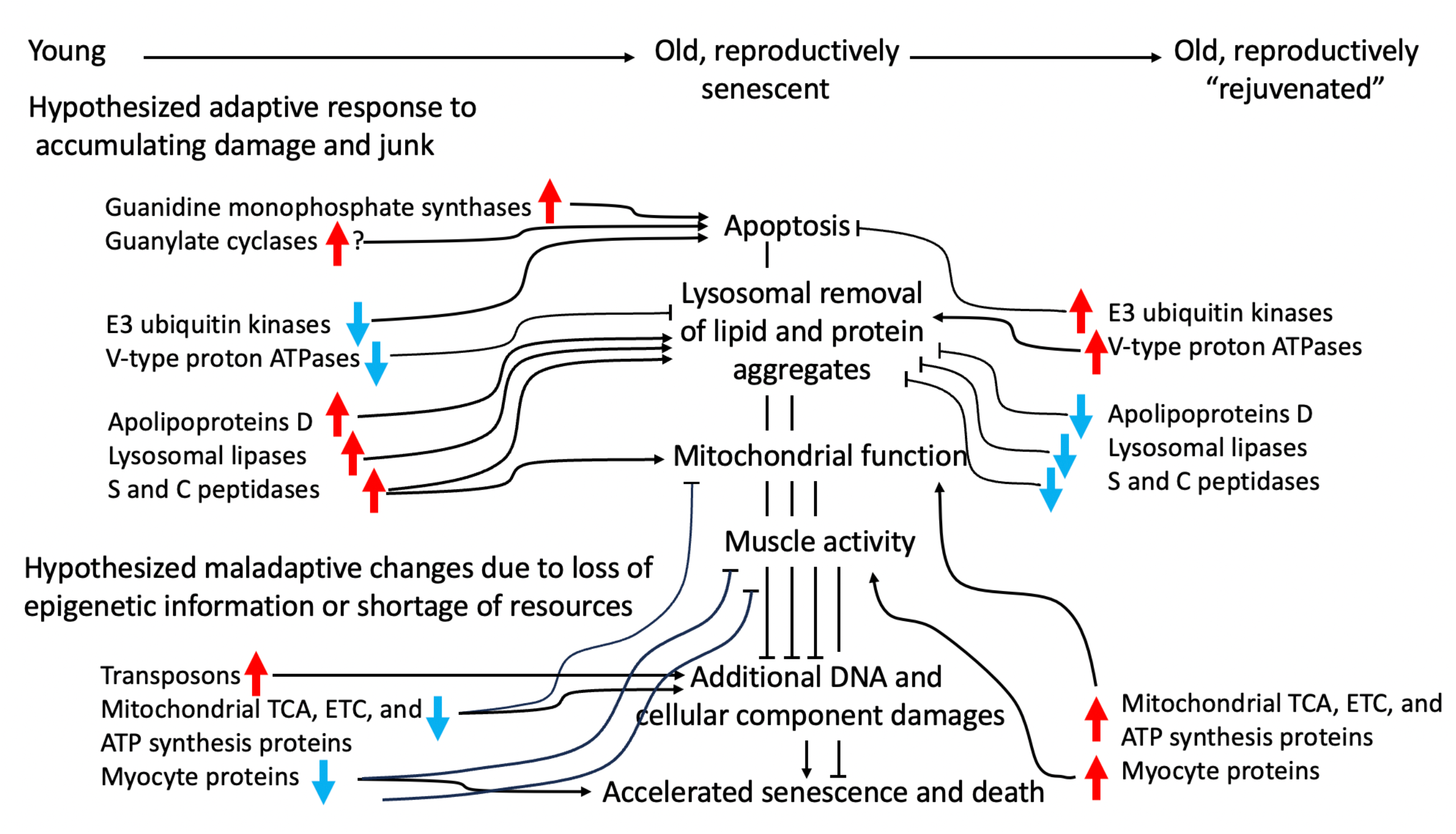
Summary of hypotheses about age-related transcriptional changes, the reversal during post-senescence reproductive events and consequences of these changes for cellular physiology and life history.

It should be noted that some of the observed transcriptional differences may be natural random variants of transcription level present constitutively, and creating a possibility for reproduction renewal. Others may be a result of active up- or down-regulation, serving as proximal causes of reproduction renewal. Others may be not necessary for it, but happen to be downstream by-products of pathways leading to such events. It is not known presently which of the observed differences allow, enable, or are the consequences of the reproduction renewal.

Four groups of paralogs stood out as those showing a strong increase in expression in old *Daphnia* without any indication of reversal in the “reproductively rejuvenated” PSRE ones: guanidine monophosphate synthases, guanylate cyclases, and two transposon-related protein families. A more diverse list of functionalities showed either the opposite trend in different paralogs, or consistent reversal or transcription activity back to “youthful” levels in the PRSE *Daphnia.* We will consider these functional groups separately.

### Differential expression between young vs. old Daphnia GMP synthases

Overexpression of GMP synthases (GMPSs) in old *Daphnia,* regardless of reproductive status, is surprising, as is the massive duplication of SMPS genes in *Daphnia* genome. It is not immediately obvious why the main function of GMPS, de novo synthesis of guanine nucleotide, should be upregulated in an old *Daphnia* in which little growth a few cell divisions occur.

However, GMPSs are multifunctional proteins with additional functions far beyond the traditional purine biosynthesis function. GMPS has been shown to play a role in deubiquitylation of regulatory proteins, most importantly of one of histones and p53, by interacting with the epigenetic silencer USP7 (van der Knaap et al. 2005). By facilitating p53 deubiquitylation the GMPS/USP7 complex results in stabilization of p53 content in cells, both in *Drosophila* and human cells (van der Knaap et al. 2005; Reddy et al. 2014). The p53 stabilization function is thought to be related to the nucleotide deficiency caused by or signaling the presence of replication or DNA-repair related stress (Bester et al. 2011; Reddy et al. 2014). Thus, elevated expression of GMPSs in old *Daphnia* may have nothing to do with purines biosynthesis, but rather with regulation of downstream targets of p53, including cell proliferation and apoptosis. Perhaps curbing the age-related accumulation of senescent or otherwise dysfunctional cells is the central target of upregulation of GMPSs. By deubiquitylation of histone H2B in *Drosophila*, GMPS is contributing to epigenetic silencing of homeotic genes by *Polycomb*. Furthermore, and, perhaps consequential for all arthropods, the GMPS/USP7 complex has been shown interact with the ecdysone (molting hormone) receptor (EcR), acting as a transcriptional corepressor of ecdysone target genes (van der Knaap et al. 2005). This is consistent with observations of molting in old-age *Daphnia,* in which molting becomes less frequent and often causes prolonged difficulties and even death. Finally, this remarkably versatile protein has also been shown to regulate accumulation of cytoplasmic triglyceride droplets in zebrafish. This function is related to de novo purines synthesis, which appears to activate ROS production, in turn upregulating the triglyceride hydrolase gene, ultimately triggering utilization of stored triglycerides (Nussbaum et al. 2013). This is consistent with reduced lipid content in very old *Daphnia* (Yampolsky et al in preparation) and with other lipid metabolism related transcriptional changes observed in this study (see below). At present it is not known which of these diverse functions are the reason for the drastic upregulation of GMPSs in aging *Daphnia,* whether this upregulation is the cause or a consequence of age-related changes, or why does *Daphnia* genome contain so many GMPS paralogs.

### Guanylate cyclases

It is perhaps not surprising, given the upregulation of GMPSs, that this family of enzymes utilizing GMPS product also showed age-dependent transcription. Four paralogs of guanylate cyclase (GC) showed up-regulation in old *Daphnia,* one of which also appearing to partly reverse this up-regulation in the PSRE females. In the case of GCs, however, the explanation for late age upregulation is readily available: the product of GC, cyclic GMP (cGMP) is a second messenger with a variety of functions most likely operating through activation of protein kinase G (PKG).

Many of the downstream pathways have long been known to regulate apoptosis. Interestingly, accumulation of cGMP can either induce (Taimor et al. 2000; Fallahian et al. 2011) or suppress (McGee et al. 1995; Wang et al. 2006) apoptosis, depending on cellular environment. Relevant to the GMPS functions discussed above, cGMP induces apoptosis under oxidative stress by activating p53 (Beyfuss & Hood 2018). Partial reversal of one of GC paralog’s expression in the PSRE females is intriguing, but at present we can only hypothesize that it may have to do with suppression of apoptosis or activation of cell proliferation in reproductive tissues. It did not escape our attention that there was a pair of paralogs showing opposite directions of age-related transcription rates are serine-threonine protein kinases, although there is no evidence that these two kinases are in fact cGMP-dependent (see below).

These results taken together suggest that there is a profound upregulation of cGMP- and p53-related apoptosis-enhancing pathways in aging *Daphnia* likely aiming at elimination senescent, damaged, or transcriptionally aberrant cells.

### Transposon-related transcripts

Another group of genes we found to consistently show higher transcription levels in aging *Daphnia* were those showing sequence similarity to with transposons-encoded genes, namely to the gag-pol polyprotein present in retroviruses and to DNA transposons’ transposase. It should be noted that there is less confidence in annotation of these proteins, as, for example, the existing genome annotation could not be confirmed by PANTHER analysis. Our assumption that these proteins are indeed transposon-encoded relies on 29% sequence identity with a retrovirus-related Pol polyprotein from transposon TNT 1-94 present in nematode *Trichinella spiralis* and 43% sequence identity with the tigger transposable element-derived protein from the bumblebee (*Bombyx mori*), respectively. Yet, assuming that these transcripts are indeed transposon-encoded, their upregulation in the old *Daphnia* is interesting. There has been recently an explosion of studies demonstrating that reactivation of transposons, retrotransposons in particular, as a consequence of erosion of epigenetic regulation that occurs with age, significantly contributes to downstream damage (Cardelli 2018; Gorbunova et al 2021; Yushkova & Moskalev 2023; Liu et al 2023). If confirmed, the increased transposon activity in aging *Daphnia* may make this organism an attractive model for the research of transposon’s role in aging.

### Differential expression between PSRE and non-PSRE old Daphnia

In nearly all cases whenever DE was observed between these categories, there was also DE between young and old, reproductively senescent *Daphnia,* with the PSRE ones partly of fully reversing these changes. It appears that, consistent with the predictions, the common themes of gene families and protein functional groups showing this pattern are the removal of lipid and protein aggregates, apoptosis, and mitochondrial functionality. Perhaps the most striking reversals has been observed in apolipoproten D paralogs (ApoD’s) and in several lysosomal lipases. Human and *Drosophila* ApoD’s have long been known to play a key role in counterbalancing oxidative stress and other damages in aging cells and, at least in *Drosophila,* increase lifespan (Sanchez et al. 2006; Walker et al. 2006; Ganfornina et al. 2008; Muffat et al. 2008; Muffat & Walker 2010; Johnson & Stolzing 2019). In flies, overexpression of either human ApoD and native *Drosophila* orthologs, *GLaz* and *Nlaz* reduces age-associated lipid peroxide accumulation and increases lifespan (Walker et al. 2006; Muffat et al. 2008), while loss of function *GLas* mutants have reduced lifespan (Sanchez et al. 2006). This regulation is paralog- and sex-specific (Ruiz et al. 2011), indicating possible connection to female reproductive function.

Likewise, what we know about lysosomal lipases and their role in aging is consistent with the above hypothesis. Overexpression of lysosomal lipases increases lifespan in *C. elegans* and this occurs through activation of autophagy (Wang et al. 2008; Lapierre et al., 2011, 2012; O’Rourke et al. 2013; Folick et al. 2015; Johnson & Stolzing 2019). Although this mechanism can operate also in germline-less worms (Lapierre et al., 2011), there is compelling evidence that it occurs through interaction between somatic maintenance and the germ line (Wang et al. 2008), again indicating a connection with reproduction function and a role reproduction-longevity trade-off.

The idea that DE of ApoD’s and lysosomal lipases in the old *Daphnia* and its reversal in PSRE ones may be interrelated, is further corroborated by the fact that the role of ApoD in ameliorating severity of the neurogenerative Niemann–Pick disease occurs through preservation of lysosomes function (Pascua-Maestro et al. 2020). Because this rescuing of lysosome function occurs through restoring pH gradient across the lysosome membrane, this may be a mechanism compensating the age-related loss of transcription of V-type proton ATPases also observed in this study. Furthermore, more recently human ApoD has been identified as not just marker or aging, but also a potential tool for achieving rejuvenation in human skin tissue (Takaya et al 2023).

With this in mind, the up-regulation of ApoD’s and lysosome lipases in aging *Daphnia* can be interpreted as the response to age-related oxidative stress operating through activation of liposomal removal of lipid aggregates, TAG hydrolysis, and autophagy. Then, the across-the board reversal of ApoDs’ and lysosomal lipases’ expression in the PSRE *Daphnia* can be interpreted in several possible ways. Are the PSRE individuals simple those who, due to random developmental variation, have not accumulated products of oxidative stress, making it both unnecessary to up-regulate the ApoD’s, and possible to resume reproduction? Or did previous overexpression of these proteins during non-reproductive phase reduced such accumulation, thus being causative for the future reproductive “rejuvenation”? Or is the anti-aging, oxidative stress-reducing activity of ApoD’s and lipases not compatible with active reproduction, requiring the PSRE *Daphnia* to bypass the anti-aging protective plasticity? If the latter, then is perhaps ApoDs’ and lysosomal lipases’ activity related to a fundamental trade-off between maintenance and reproduction? It is not immediately obvious what experiments might help differentiate between these possibilities. A knock-out study might, but it would be necessary to knock out several paralogs in each gene family.

Similar reasoning, perhaps related to both apoptosis and lysosomal removal of protein aggregates applies to the same pattern observed in serine and cysteine proteases: upregulation in aging Daphnia reversed in the PSRE ones. We know that various serine proteases participate in a variety of aging-related aggregate removal (Yamin et al 2009), apoptosis (Madeo et al. 2004; Fahrenkrog 2011), and mitochondrial homeostasis (Kang et al. 2013; see below) pathways in yeast and mammals. However, the extreme diversity of functions of serine- and cysteine proteases makes it difficult to formulate specific hypotheses.

It is more difficult to interpret the transcriptional pattern observe in V-type proton ATPases, whose central function is maintaining acidic pH in lysosomes, critical for lysosomes’ lipid and protein hydrolysis function (Beyenbach & Wieczorek 2006). On the one hand partial restoration of their RNA abundancies in the PSRE *Daphnia* is to be expected (cf. Bohnert & Kenyon 2017). But the reduced transcription in senescent *Daphnia* does not fit the above damage-control reasoning applied to ApoD’s and lipases. Perhaps here we observe the situation in which age-related transcriptional changes are caused not by mobilization of protective mechanisms, but by their decline. Either maintaining low pH in lysosomes becomes too expensive (perhaps due to reduced ATP production, see below) or unnecessary due to reduced reproduction, if the role of these ATPases is similar to that described by Bohnert and Kenyon (2017), i.e. crucial for mainlining rejuvenated germline.

The other gene family repeatedly showing reduction in RNAs abundancies in old *Daphnia* that is partly reversed in the PSRE individuals were the E3 ubiquitin-protein ligases. Specific hypotheses are difficult to make due to a great variety of substrates for these kinases, but various ubiquitin-protein ligases have been implemented in protein degradation (Hershko 2005), as well as regulation of autophagy, including mitophagy (Hattori and Mizuno 2017), and apoptosis by targeting p53 for degradation (Vazquez et al. 2008; Pawge & Khatik 2021). Thus, downregulation of ubiquitin-protein ligases in old *Daphnia* is consistent with elevated apoptosis, a trend partly reversed in the PSRE individuals.

Finally, and perhaps most predictably, we observed full or partial restoration of transcription levels to those observed in young individuals in several key proteins responsible for mitochondrial function and muscle activity. Both electron-transport chain (components of complex I and II, but not complex III or IV) and ATP synthase components showed this pattern. The largest age-related difference, almost 60-fold, was observed in the abundance of ATP synthase epsilon subunit, much of which was reversed in the PSRE individuals. In *Drosophila* e-ATP synthase subunit, in addition to structural role as part of ATP synthase rotor, is critical for proper mitosis in the embryo and serves as a ligand for the aging regulator Methuselah (Kidd et al. 2005), playing a central role in fat body nutrient level signaling (Delanoue et al. 2016). It is not know whether the mitochondrial ATP synthase or nutrient signaling functions of this protein is critical to the observed age-related DE. Given that other ATP subunits did not show DE, the latter function may be of a greater importance in *Daphnia* aging and reproductive rejuvenation. It should be noted that the observed age-related reduction of ETC components expression occurs without any weight-adjusted changes in oxygen consumption in aging *Daphnia* (Lowman & Yampolsky 2023). This is consistent with no DE in complex IV of the ETC and may indicate that aging mitochondria are less efficient in maintaining proton gradient per mole of oxygen consumed. On the other hand, downregulation of several myocyte-specific proteins is consistent with the observed reduction of locomotory and filter-feeding activity in aging *Daphnia* (Anderson et al. 2022; Pearson et al., in preparation). Because *Daphnia,* as any other zooplankton filter-feeding organisms, has to constantly swim to stay in the water column and filter water to obtain food, reduced muscular activity in nature is likely to lead to mortality.

One other pattern of transcriptional changes associated with aging and PSREs we have observed was the opposite direction of DE changes in different paralogs, suggestive of possible aging-related specialization, and, in some cases, possibly paralog retention by subfunctionalization. There were several gene families showing this pattern, some of which with functionality without any a priori connection to aging or rejuvenation. These include ATP-dependent RNA helicases, chorion peroxidases, cuticle proteins, solute carriers, T-complex proteins, and zinc finger proteins. Two other functional groups, namely G-protein coupled receptors and serine/threonine-protein kinases, have known aging-related functionalities, but, again, the large number of paralogs within these groups in most genomes and the great diversity of their functions make it difficult to formulate specific hypotheses about the roles of these particular paralogs in aging and rejuvenation.

## Conclusions

Post-senescence reproductive evens in *Daphnia* are not followed by elevated mortality as predicted by the terminal investment hypothesis. Rather, they represent a transcriptionally unique state in which transcription of several gene families and functional groups of genes showing age-related decline or increase in transcription is reversed to the “youthful” levels.

Functionally, this pattern is observed in genes implicated in lipid metabolism, lysosomal functions, removal of lipid and protein aggregates, apoptosis, autophagy, and mitochondrial and muscle functions.

## Acknowledgments

We are grateful to Cora Anderson and Morad Malek for laboratory assistance and to Marc Kirschner for useful discussions. RNA-seq library preparation and initial analysis were performed by the Functional Genomics Core, COBRE Center for Targeted Therapeutics at the University of South Carolina. Funding: Impetus Foundation grant to LYY.

## Data Availability

All life-history data necessary to recreate the results reported here are available from Supplementary Data file. Raw and mapped reads from the RNAseq experiment are available from NCBI, BioProject PRJNA1098345; SRA Accession numbers SRR28604180 - SRR28604191; processed data from NCBI GEO GSE263788, Accession numbers GSM8199167 - GSM8199178.

