## Supplementary Tables and Figures for "Post-senescence reproductive rebound in *Daphnia* associated with reversal of age-related transcriptional changes"

A

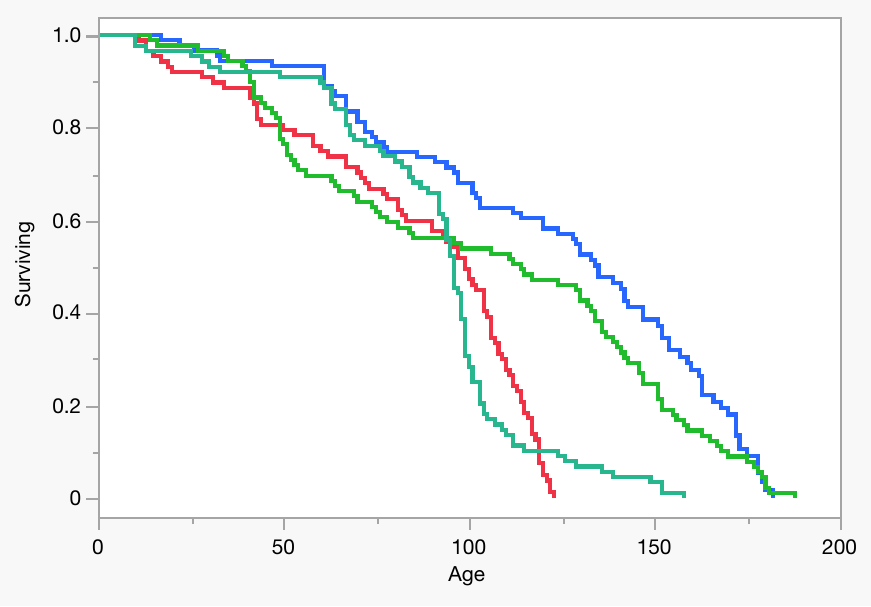

p < 0.0001

p < 0.0001

p < 0.061

p < 0.011

p < 0.0001

p < 0.054

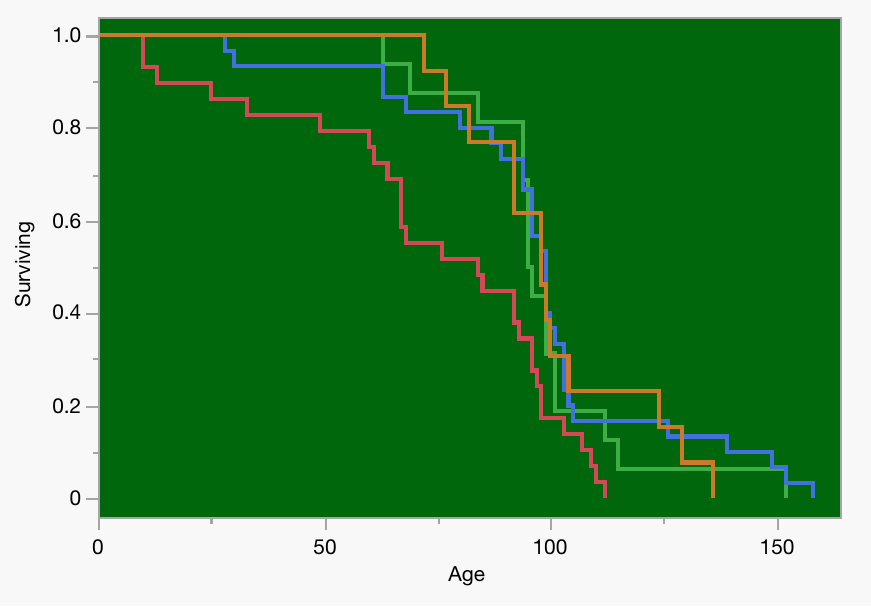
B	C	D	E

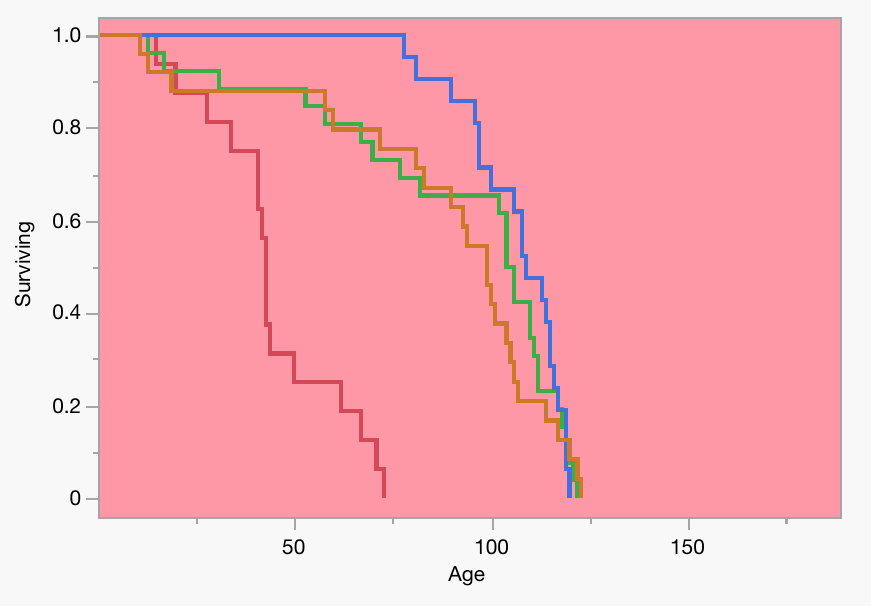

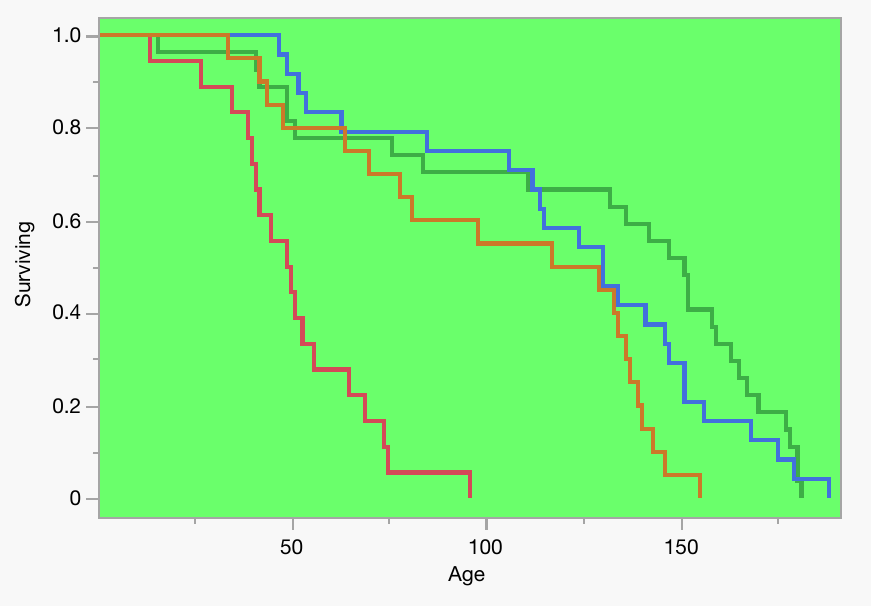

p < 0.0001

p < 0.0001

p < 0.011

p < 0.0044

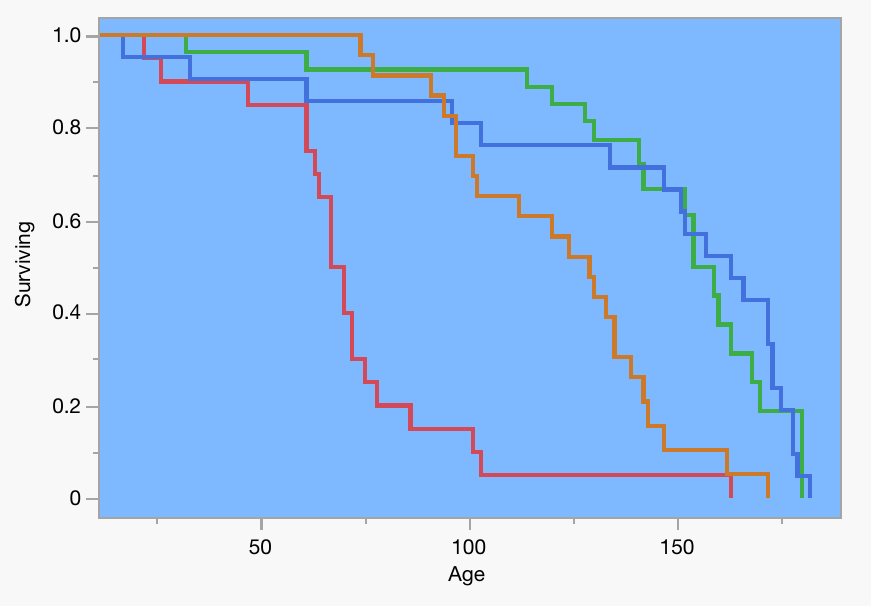

FI	IL	HU	GB

p < 0.0001

p < 0.0001

p < 0.0001

p < 0.0001

Fig. S1. Overall survival curves in two lifespan experiments at different temperatures and medium compositions. A: 4 clones combined. B – E: 4 clones at different conditions (labels correspond to the first 2 letters of clones’ ID. Colors of curves on A and backgrounds on B - E as on main text Fig.X: blue, Experiment 2, ADaM medium, 20 ^o^C; red, Experiment 2, ADaM medium, 24 ^o^C; light green Experiment 2, COMBO medium, 20 ^o^C; dark green Experiment 1, COMBO medium, 20 ^o^C. See Supplementary Table XXX for full analysis. In each contrast the top p-value reported is from a Log-Rank test (more sensitive to late-age mortality) and the bottom p-value is from the Wilcoxon test (more sensitive to early mortality; Kalbfleisch and Prentice [1980](https://www.jmp.com/support/help/en/17.0/jmp/references-10.shtml#ww127422)).

The Log-Rank test places more weight on larger survival times and is more useful when the ratio of hazard functions in the groups being compared is approximately constant. The hazard function is the instantaneous failure rate at a given time. It is also called the *mortality rate* or *force of mortality*.

The Wilcoxon test places more weight on early survival times and is the optimum rank test if the error distribution is logistic. See Kalbfleisch and Prentice ([1980](https://www.jmp.com/support/help/en/17.0/jmp/references-10.shtml#ww127422)).

Kalbfleisch, J. D., and Prentice, R. L. (1980). *The Statistical Analysis of Failure Time Data*. New York: John Wiley & Sons.

Mean clutch size (Total offspring)

| Age class, days | Non-PSRE | PSRE |
| --- | --- | --- |
| 0 | 4.02586 | 5.28571 |
| 20 | 4.35594 | 5.03005 |
| 40 | 4.46758 | 4.89089 |
| 60 | 4.65504 | 5.31486 |

Fig. S2. Mean fecundity (clutch size, total offspring) in pre-senescent *Daphnia* that later in life did or did not demonstrate the PSREs. See Supplementary Table S3 for detailed analysis.

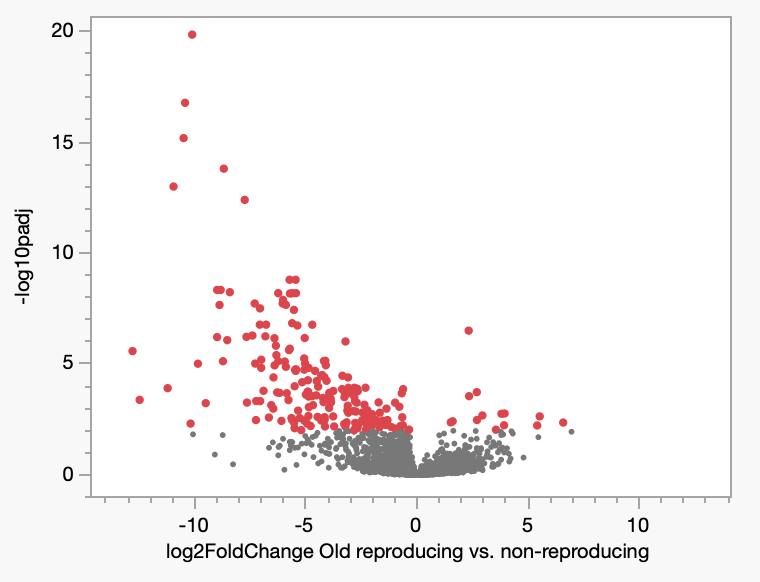

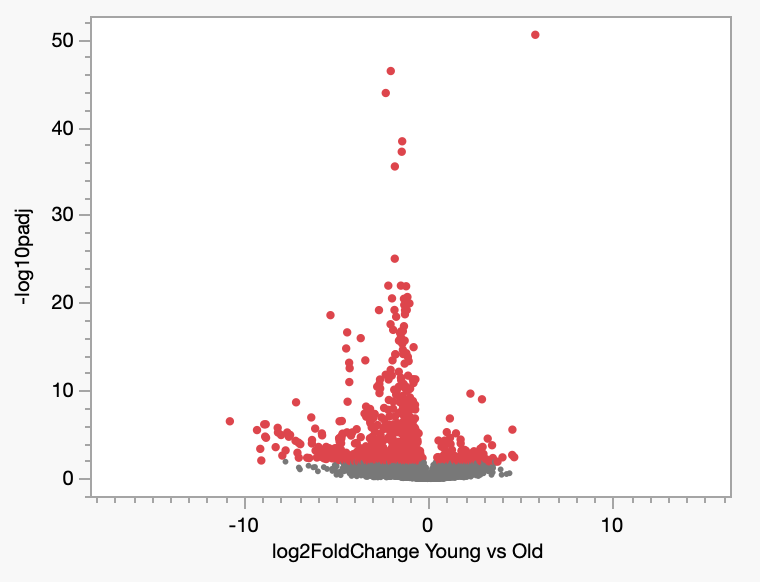

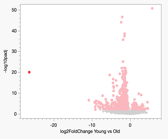

Fig. S3. Volcano plots showing genes differentially expressed in young vs. old Daphnia (left) and PSRE vs. non-PSRE old Daphnia. Transcripts with P_adj_<0.01 shown in red. Positive logFC values correspond to higher expression in young (left panel) and in PSRE old individuals. Insert: zoom out to show an ApoD paralog expressed only in old Daphnia.

GMPSs

DCq

|  | Young | PSRE | non-PSRE |
| --- | --- | --- | --- |
| GMPS_038F5.143 | 10.97 | 10.6225 | 10.4983 |
| GMPS_00F30.73 | 11.4038 | 10.1113 | 10.5617 |
| GMPS_001F48.110 | 4.29125 | 2.84375 | 2.65833 |

ApoDs

DCq

|  | Young | PSRE | non-PSRE |
| --- | --- | --- | --- |
| ApoD_014F14.88 | 7.53375 | 8.075 | 6.54833 |
| ApoD_014F14.90 | 12.95 | 11.1325 | 7.64333 |
| ApoD_014F14.91 | 8.7325 | 8.8025 | 6.09333 |
| ApoD_014F14.86 | 14.24 | 12.7217 | 3.86833 |

GC

DCq

|  | Young | PSRE | non-PSRE |
| --- | --- | --- | --- |
| GC_065F2.33 | 5.92875 | 4.77125 | 4.7 |

*

*

Fig. S4. qPCR confirmation of differences in select transcript abundances. ΔC_q_: cycles to threshold, control gene (*Xbp*) values subtracted. (higher values correspond to lower initial template abundance). Colors as on main text Figs 6 – 8. Horizontal lines show significance of pairwise difference between categories by Dunnett’s test (* p<0.05; ** p<0.01; *** p<0.001). Bars are SE reflecting differences between biological replicates. Full statistics in Supplementary Table S7.

Table S0. Location and habitat type of origin of Daphnia clones used. First two letters of Clone ID are used to identify clones throughout the paper.

*Anderson et al. 2022

| **CloneID** | **Type of Habitat** | **Latitude** | **Longitude** | **Median lifespan, days * (95% CI)** |
| --- | --- | --- | --- | --- |
| FI-FSP1-16-2 | Intermittent (summer rock pool) | 60° 10' 40" N | 25° 47' 45" E | 36 (22-42) |
| GB-EL75-69 | Permanent (year-round pond) | 51° 30' 26" N | -0° 7' 39" E | 64 (62-68) |
| HU-K-6 | Semi-permanent (saline lake) | 46° 47' 33" N | 19° 10' 54" E | 72 (67-77) |
| IL-MI-8 | Intermittent (Mediterranean summer-dry pond) | 31° 46' 40" N | 35° 13' 14" E | 54 (46-57) |

Table S1. Proportional hazards analysis (Wald’s test) of survival in two lifespan experiments at different temperatures and medium compositions.

| Source | DF | Chi^2^ | Prob>Chi^2^ |
| --- | --- | --- | --- |
| Experiment | 1 | 5.41 | 0.02 |
| Treatment | 3 | 76.13 | <.0001 |
| Clone | 3 | 179.48 | <.0001 |
| Clone*Treatment | 12 | 50.15 | <.0001 |

Table S2. Nominal logistic model of the effects of body size at maturity on late-life PSREs. Only individuals with removal date (date dead or censored) beyond the upper limit of each age class included into the data subset. Analysis across both experiments and four clones (including the FI clone that showed no PRSE in Experiment 2).

| Source | DF | Chi^2^ | Prob> Chi^2^ |
| --- | --- | --- | --- |
| Treatment | 3 | 4.06 | 0.25 |
| Clone | 3 | 311.62 | <0.0001 |
| Treatment*Clone | 8 | 12.98 | 0.11 |
| L, mm | 1 | 6.0E-06 | 0.998 |
| Treatment*L, mm | 3 | 4.01 | 0.26 |
| Clone*L, mm | 3 | 112.38 | <0.0001 |
| Treatment*Clone*L, mm | 7 | 19.76 | 0.0061 |

Table S3. Nominal logistic model of the effects of pre-senescence fecundity on late-life PSREs occurrence with differences among treatments and clones accounted for. Only individuals with removal date (date dead or censored) beyond the upper limit of each age class included into the data subset. Analysis across both experiments and four clones (including the FI clone that showed no PRSE in Experiment 2).

| Age class | | 0 - 10 | | 10 - 30 | | 30 - 50 | | | 50 - 70 | | |
| --- | --- | --- | --- | --- | --- | --- | --- | --- | --- | --- | --- |
| Source | DF | Chi^2^ | Prob>Chi^2^ | Chi^2^ | Prob> Chi^2^ | | Chi^2^ | Prob>Chi^2^ | | Chi^2^ | Prob>Chi^2^ |
| Mean clutch size | 1 | 1.189 | 0.28 | 4.011 | 0.045 | | 0.072 | 0.79 | | 0.989 | 0.32 |
| Treatment | 3 | 0.544 | 0.91 | 2.207 | 0.53 | | 10.798 | 0.013 | | 4.181 | 0.24 |
| Clone | 3 | 5.287 | 0.15 | 52.104 | <.0001 | | 36.604 | <.0001 | | 25.975 | <.0001 |
| Treatment*Clone | 8 | 8.019 | 0.43 | 10.794 | 0.29 | | 13.930 | 0.12 | | 11.906 | 0.22 |

Table S4. Results of GO and pathway enrichment analysis (terms with FET FDR<0.05, filtered down to the narrowest term). N_DE_, N_ref_, N_exp_: counts of genes in the DE list, in the reference and expected count, respectively. GO or pathway terms that are enriched by enrichment of the same set of differentially expressed genes are grouped together. When multiple terms show enrichment due to enrichment of the same gene family gene count shown only for the term with the lowest FDR.

| GO or Pathway terms | ID | FET FDR | N_DE_ | N_ref_ | N_exp_ | Underlying enriched gene family / function |
| --- | --- | --- | --- | --- | --- | --- |
| Enriched among genes **up-**regulated in **old,** non-reproducing *Daphnia* relative to young *Daphnia* | | | | | | |
| Biological process, molecular function of pathway | | | | | | |
| De novo purine biosynthesis^a^ | P02738 | 7.6E-11 | 25 | 170 | 4.49 | GMP synthases |
| purine ribonucleotide biosynthetic process | GO:0009152 | 1.7E-09 |  |  |  |  |
| ligase activity | GO:0016874 | 7.5E-08 |  |  |  |  |
| apolipoprotein^b^ | PC00052 | 0.0008 | 7 | 34 | 0.76 | Apolipoprotein D’s |
| serine-type peptidase activity^c^ | GO:0008236 | 0.0016 | 9 | 66 | 1.52 | Ser endopeptidases |
| cysteine-type endopeptidase activity | GO:0004197 | 0.0153 | 7 | 53 | 1.22 | Cys endopeptidases / cathepsins |
| Cellular localization | | | | | | |
| extracellular space^d^ | GO:0005615 | 5.4E-07 | 28 | 329 | 7.57 | Proteases, cuticular proteins, collagens |
| cytosol^a^ | GO:0005829 | 0.0383 | 25 | 521 | 12.0 | GMP synthases |
| Enriched among genes **down**-regulated in **old**, non-reproducing *Daphnia* relative to young *Daphnia* | | | | | | |
| ribosomal protein | PC00202 | 4.2E-10 | 38 | 176 | 9.56 | Ribosomal proteins |
| large ribosomal subunit | GO:0015934 | 3.4E-05 |  |  |  |  |
| small ribosomal subunit | GO:0015935 | 7.9E-05 |  |  |  |  |
| translational elongation | GO:0006414 | 5.9E-05 | 29 | 184 | 10.0 | Ribosomal proteins, elongation factors, AARS’es |
| amide biosynthetic process | GO:0043604 | 0.0008 |  |  |  |  |
| RNA processing | GO:0006396 | 0.0070 |  |  |  |  |
| translational initiation | GO:0006413 | 0.037 |  |  |  | Translation initiation factors |
| Cellular localization | | | | | | |
| Cytosol | GO:0005829 | 9.8E-04 | 25 | 521 | 12.0 | Various |
| t-UTP complex | GO:0034455 | 0.0059 | 10 | 40 | 2.2 | Various |
| Depleted among genes **down**-regulated in **old**, non-reproducing Daphnia relative to young Daphnia | | | | | | |
| viral or transposable element protein | PC00237 | 1.4E-11 | 0 | 629 | 32.2 | None |
| Enriched among genes up-regulated in old, non-reproducing *Daphnia* relative to old, reproducing (PSRE) *Daphnia* | | | | | | |
| Biological process, molecular function of pathway | | | | | | |
| apolipoprotein^b^ | PC00052 | 0.0002 | 6 | 34 | 0.21 | Apolipoprotein D’s |
| response to oxidative stress | GO:0006979 | 0.0003 |  |  |  |  |
| transfer/carrier protein | PC00219 | 0.0035 |  |  |  |  |
| Lipase | PC00143 | 0.0004 | 6 | 47 | 0.29 | Lysosomal lipase-related |
| lipid metabolic process | GO:0006629 | 0.0018 |  |  |  |  |
| serine-type endopeptidase activity^c^ | GO:0004252 | 0.0190 | 5 | 65 | 0.40 | Ser endopeptidases |
| Cellular localization | | | | | | |
| extracellular space^e^ | GO:0005615 | 0.0010 | 12 | 325 | 2.05 | Proteases, cuticular proteins |

^a^ enriched in more than one DE category due to the same differentially expressed genes.

^b^ enriched in more than one DE category due to the same differentially expressed genes.

^c^ enriched in more than one DE category due to the same differentially expressed genes.

^e^ differentially expressed genes are a subset of those in ^d^.

Table S5. Individual genes or gene families (not contributing to GO enrichment, Table S4) that are up-regulated in old non-reproducing Daphnia relative to young ones (DEseq2 adjusted P<0.0001).

| paralogs family | N paralogs | | Max(-log10(padj)) |
| --- | --- | --- | --- |
| Regulator of G-protein signaling | 1 | 11.4 | |
| 25-hydroxyvitamin D-1 alpha hydroxylase, mitochondrial | 1 | 11.1 | |
| Vitamin D 25-hydroxylase | 2 | 11.1 | |
| cpg binding protein | 1 | 10.1 | |
| BAI1-associated protein | 1 | 9.5 | |
| phenylalanine-4-hydroxylase | 1 | 9.2 | |
| Cytochrome P450 315a1, mitochondrial | 1 | 8.4 | |
| Sugar transporter ERD6 | 3 | 8.3 | |
| Carbonic anhydrase | 1 | 8 | |
| Transcription factor | 1 | 7.5 | |
| Bestrophin | 2 | 7.3 | |
| ATP-binding cassette containing protein | 4 | 6.7 | |
| Voltage-gated sodium channel subunit alpha NaV1.7 | 1 | 6.6 | |
| Pol protein | 5 | 6.3 | |
| Receptor-type tyrosine-protein kinase FLT3 | 1 | 5.8 | |
| transposases | 3 | 5.7 | |
| Kinesin-associated protein | 1 | 5.7 | |
| Cell division protein ftsj | 4 | 5.5 | |
| 63 kDa sperm flagellar membrane protein | 1 | 5.4 | |
| Elongation of very long chain fatty acids protein | 2 | 5.4 | |
| TE: Retrovirus-related Pol polyprotein from | 1 | 5.3 | |
| Serine/threonine-protein kinases | 1 | 4.9 | |
| Opioid-binding protein/cell adhesion molecule | 1 | 4.8 | |
| Galactose-3-O-sulfotransferase | 2 | 4.8 | |
| Calcium-activated chloride channel regulator | 1 | 4.7 | |
| Helicase senataxin | 1 | 4.5 | |
| Nuclear receptor subfamily 6 group A member 1 | 1 | 4.4 | |
| U4/U6.U5 tri-snRNP-associated protein | 1 | 4.4 | |
| Sorting nexin-5 | 1 | 4.3 | |
| NADPH oxidase | 1 | 4.3 | |
| BarH 1 homeobox protein | 1 | 4.3 | |
| Neurexin IV | 1 | 4.3 | |
| Brain chitinase and chia | 1 | 4.3 | |
| DMRT2 protein | 1 | 4.1 | |
| Glioma pathogenesis-related protein | 1 | 4.1 | |
| Lactosylceramide 4-alpha-galactosyltransferase | 1 | 4.1 | |
| Melanization protease 1, | 1 | 4.1 | |
| BTB/POZ domain-containing protein KCTD21 | 1 | 4 | |
| Ribonuclease P protein subunit p40 | 1 | 4 | |
| Glucosyl/glucuronosyl transferases | 2 | 4 | |
| myosin xv | 1 | 3.9 | |
| Cral/trio domain-containing protein | 1 | 3.9 | |
| udp-galactose:n-acetylgalactosamine-alpha-r beta galactosyltransferase | 1 | 3.9 | |
| h3 histone | 1 | 3.8 | |
| calpa | 1 | 3.8 | |
| Synaptotagmin-17 | 1 | 3.8 | |
| Pheromone and odorant receptor | 1 | 3.8 | |
| Beta-1,3-galactosyltransferase 2 | 1 | 3.7 | |
| Ankyrin repeat and SOCS box protein 2 | 1 | 3.6 | |
| AT14086p | 1 | 3.5 | |
| Jmjc domain-containing histone demethylation protein | 1 | 3.5 | |
| Estradiol 17-beta-dehydrogenase 12-B | 1 | 3.5 | |
| Tubulin polyglutamylase TTLL7 | 1 | 3.4 | |
| RT_nLTR_like, RT_nLTR: Non-LTR (long terminal repeat) retrotransposon and | 1 | 3.4 | |
| Homeobox protein Hox-A2 | 1 | 3.4 | |
| 1-acyl-sn-glycerol-3-phosphate acyltransferase delta | 1 | 3.4 | |
| MAP kinase-activating death domain protein | 1 | 3.3 | |
| Carboxypeptidase A4 | 1 | 3.3 | |
| sialin | 1 | 3.3 | |
| Angiotensin-converting enzyme | 1 | 3.3 | |
| Neuroglobin | 1 | 3.2 | |
| Cadherin-13 | 1 | 3.2 | |
| Plasminogen | 1 | 3.2 | |
| Leucine zipper tumor suppressor 1, putative | 1 | 3.2 | |
| Fatty acyl-CoA reductase | 1 | 3.2 | |
| Quinone oxidoreductase PIG3 | 1 | 3.2 | |
| polyprotein of retroviral origin | 1 | 3.1 | |
| cGMP-inhibited 3',5'-cyclic phosphodiesterase B | 1 | 3.1 | |
| Complement C1r subcomponent | 1 | 3.1 | |
| Low-density lipoprotein receptor-related protein | 1 | 3.1 | |
| MDS1 and EVI1 complex locus protein EVI1 | 1 | 3 | |
| Carboxylesterase 5A | 1 | 3 | |
| Enhancer of split mgamma protein | 1 | 3 | |
| Integral membrane protein GPR155 | 1 | 3 | |
| Menin | 1 | 3 | |

Table S6. Individual genes or gene families (not contributing to GO enrichment, Table S4) that are down-regulated in old non-reproducing Daphnia relative to young ones (DEseq2 adjusted P<0.0001).

| paralogs family | N paralogs | Max(-log10(padj)) |
| --- | --- | --- |
| Glutamine gamma-glutamyltransferase K | 2 | 8.3 |
| Ionotropic glutamate receptor subunit ia | 1 | 8.2 |
| Troponin T, fast skeletal muscle | 1 | 6.1 |
| E3 ubiquitin protein ligases | 5 | 5.6 |
| Inactive serine/threonine-protein kinase TEX14 | 1 | 5.4 |
| Platelet-activating factor acetylhydrolase | 1 | 5.2 |
| Fibrillin-1 | 3 | 5.1 |
| Defense protein l(2)34Fc | 1 | 5.1 |
| Tropomyosin alpha-1 chain | 2 | 4.8 |
| Suppressor of cytokine signaling 6 | 1 | 4.7 |
| c-type lectin ctl - mannose binding. | 1 | 4.7 |
| 5-aminolevulinate synthase, nonspecific, mitochondrial | 1 | 4.4 |
| Retinol dehydrogenase | 2 | 4.3 |
| fs1m3 | 1 | 4.3 |
| Calcineurin B ous protein | 1 | 4.2 |
| ATP synthase subunit epsilon, mitochondrial | 1 | 4.2 |
| Bipolar kinesin KRP-130 | 1 | 4.1 |
| Titin | 1 | 4 |
| Myosin light chain | 1 | 3.9 |
| RNA-binding protein | 1 | 3.9 |
| Coiled-coil domain-containing protein 43 | 1 | 3.9 |
| Protein goliath | 1 | 3.9 |
| Erlin-1 endoplasmic reticulum lipid raft-associated protein | 1 | 3.8 |
| Double-strand-break repair protein rad21 | 1 | 3.8 |
| Proliferation-associated protein 2G4 | 1 | 3.8 |
| Actin-related protein | 1 | 3.8 |
| Microtubule-associated protein | 1 | 3.7 |
| Probable cytochrome P450, mitochondrial | 2 | 3.7 |
| Phospholipase B lamina ancestor | 1 | 3.7 |
| Basement membrane-specific heparan sulfate proteoglycan core protein | 1 | 3.7 |
| Profilin-4 | 1 | 3.6 |
| Squamous cell carcinoma antigen recognized by t-cells 3 sart-3 msart-3 tumor-rejection antigen sart3 | 1 | 3.5 |
| Ribosome biogenesis protein BRX1 homolog | 1 | 3.5 |
| Globin | 1 | 3.5 |
| Hsp90 co-chaperone Cdc37 | 1 | 3.5 |
| DNA topoisomerase I, mitochondrial protein | 1 | 3.5 |
| Nucleolar RNA helicase 2 | 1 | 3.5 |
| Methionine aminopeptidase | 1 | 3.5 |
| Nucleosome assembly protein 1 4 | 1 | 3.4 |
| Thymocyte selection-associated high mobility group box protein TOX | 1 | 3.4 |
| Dihydrolipoyllysine-residue acetyltransferase component of pyruvate dehydrogenase complex | 1 | 3.4 |
| UDP-N-acetylglucosamine--peptide N-acetylglucosaminyltransferase 110 kDa subunit | 1 | 3.4 |
| Mpv17 | 1 | 3.4 |
| Nucleolar NOP5 protein | 2 | 3.4 |
| Midasin | 1 | 3.4 |
| Cutaneous T-cell lymphoma-associated antigen 5 | 1 | 3.3 |
| Vitellogenin-1 precursor | 1 | 3.3 |
| Importin-4 | 1 | 3.3 |
| Basic leucine zipper and W2 domain-containing protein 1-A | 1 | 3.3 |
| Myosin light chain kinase, smooth muscle | 2 | 3.3 |
| UDP-GlcNAc:betaGal beta-1,3-N-acetylglucosaminyltransferase | 1 | 3.3 |
| Krueppel factor 15 | 1 | 3.3 |
| SWI/SNF-related matrix-associated actin-dependent regulator of chromatin subfamily A member 5 | 1 | 3.2 |
| Integrin alpha-6 | 1 | 3.2 |
| plasminogen activator inhibitor 1 rna-binding protein | 1 | 3.2 |
| Transmembrane and TPR repeat-containing protein | 1 | 3.2 |
| Microsomal triglyceride transfer protein large subunit | 1 | 3.2 |
| Trafficking protein particle complex subunit | 1 | 3.1 |
| Hydroxysteroid dehydrogenase protein 2 | 1 | 3.1 |
| Nicotinic acetylcholine receptor subunit alpha | 1 | 3.1 |
| Spire | 1 | 3.1 |
| 3-methyl-2-oxobutanoate dehydrogenase [lipoamide] kinase | 1 | 3.1 |
| 60 kDa heat shock protein homolog 1, mitochondrial | 1 | 3.1 |
| Anaphase-promoting complex subunit | 1 | 3.1 |
| Splicing factor 3A subunit | 1 | 3.1 |
| Patched domain-containing | 1 | 3.1 |
| Mitochondrial inner membrane protein | 1 | 3 |
| Muscle, skeletal receptor tyrosine protein kinase | 1 | 3 |
| Dendritic cell-specific transmembrane protein | 1 | 3 |
| Maltase-glucoamylase, intestinal | 1 | 3 |

Table S7. Individual gene families (not contributing to GO enrichment, Table S4) in which some paralogs are up- and others down-regulated in old non-reproducing Daphnia relative to young ones (DEseq2 adjusted P<0.0001).

| paralogs family | N paralogs UP | N paralogs DOWN | Max(-log10(padj))  UP | Max(-log10(padj)) DOWN |
| --- | --- | --- | --- | --- |
| Chorion peroxidase precursor | 2 | 1 | 14.4 | 8.2 |
| Solute carriers | 4 | 2 | 5.4 | 3.7 |
| ATP-dependent RNA helicases | 1 | 1 | 4.2 | 4.3 |
| Cuticle proteins | 4 | 1 | 4.3 | 4.4 |
| T-complex protein 1 subunits | 1 | 1 | 3.9 | 4.1 |
| G-protein coupled receptor | 1 | 1 | 3.9 | 3.3 |
| Serine threonine-protein kinases | 1 | 1 | 3.9 | 3.7 |
| Zinc finger protein | 3 | 1 | 3.6 | 9.1 |

Supplementary Table S7. qPCR confirmation of differences in select transcript abundances. Tw-way (for each gene family) and one-way (for individual genes) ANOVAs with C_q_ as the response and age/reproductive class (young / old, PSRE / old, non-PSRE) as the main effect. Xbp used as a control. See Supplementary Fig. S4 for Tukey test results where appropriate.

| Source | DF | Sum of Squares | F | | | Prob > F |
| --- | --- | --- | --- | --- | --- | --- |
| Transcripts: GMPS’s | | | | |  |  |
| Gene | 2 | 397.53 | 210.31 | <0.0001 | | |
| AgeClass | 2 | 7.50 | 3.97 | 0.033 | | |
| Gene*AgeClass | 4 | 2.07 | 0.55 | 0.70 | | |
| Error | 24 | 22.68 |  |  | | |
| Transcripts: ApoD’s | | | | | | |
| Gene | 3 | 82.97 | 4.72 | 0.0081 | | |
| AgeClass | 2 | 175.05 | 14.95 | <0.0001 | | |
| Gene*AgeClass | 6 | 90.58 | 2.58 | 0.039 | | |
| Error | 30 | 175.6 |  |  | | |
| Transcript: GC | | | | |  |  |
| AgeClass | 2 | 3.60 | 5.08 | 0.038 | | |
| Error | 8 | 2.84 |  |  | | |
